# Identification of candidate virulence loci in *Striga hermonthica*, a devastating parasite of African cereal crops

**DOI:** 10.1101/2022.01.13.476148

**Authors:** Suo Qiu, James M. Bradley, Peijun Zhang, Roy Chaudhuri, Mark Blaxter, Roger K. Butlin, Julie D. Scholes

## Abstract

- Parasites have evolved proteins, Virulence Factors (VFs), that facilitate plant colonization, yet VFs mediating parasitic plant-host interactions are poorly understood. *Striga hermonthica* is an obligate, root-parasitic plant of cereal hosts in sub-Saharan Africa, causing devastating yield losses. Understanding the molecular nature and allelic variation of VFs in *S. hermonthica* is essential for breeding resistance and delaying the evolution of parasite virulence.
- We assembled the *S. hermonthica* genome and identified secreted proteins by *in silico* prediction. Pooled sequencing of parasites growing on a susceptible and a strongly resistant rice host allowed us to scan for loci where selection imposed by the resistant host had elevated the frequency of alleles contributing to successful colonisation.
- Thirty-eight putatively secreted VFs had extremely different allele frequencies with functions including host cell wall modification, protease inhibitors, oxidoreductase and kinase activities. These candidate loci had significantly higher Tajima’s D than the genomic background, consistent with balancing selection.
- Our results reveal diverse strategies used by *S. hermonthica* to overcome different layers of host resistance. Understanding the maintenance of variation at virulence loci by balancing selection will be critical to managing the evolution of virulence as a part of a sustainable control strategy.

## Introduction

Plants are constantly challenged by parasites from across all kingdoms of life (Win *et al*., 2012; Mitsumasu *et al*., 2015). As a consequence, they have evolved sophisticated surveillance systems to detect and protect themselves against parasite invasion (Cook *et al*., 2015; Wu *et al*., 2018; Kanyuka *et al*., 2019). In turn, plant parasites have evolved suites of proteins, miRNAs, or other molecules which are delivered into host plants to facilitate colonisation (Virulence Factors (VFs)) (Win *et al*., 2012; Giraldo *et al*., 2013; Zheng *et al*., 2013; Mitsumasu *et al*., 2015). These VFs are pivotal in determining the outcome of a parasite-plant interaction. Despite substantial advances in understanding the identity and mode of action of VFs in plant interactions with fungal, bacterial and nematode parasites (Win *et al*., 2012; Giraldo *et al*., 2013; Zheng *et al*., 2013) much less is known about VFs mediating parasitic plant interactions with their plant hosts (Westwood *et al*., 2010, 2012; Timko *et al*., 2012). Parasitic plants occur in almost all terrestrial habitats and have evolved independently at least 12 times (Kuijt 1969; Westwood *et al*., 2010; Clarke *et al*., 2019). Regardless of evolutionary origin, parasitic plants possess a multicellular organ called the ‘haustorium’, through which direct structural and physiological connections are formed with their host plant (Westwood *et al*., 2010; Yoshida *et al*., 2016). This allows them to abstract water, organic and inorganic nutrients. In addition, the haustorium is increasingly recognised to play a role in host manipulation, through the movement of parasite-derived proteins, miRNAs and other small molecules into the host plant (Aly *et al*., 2011; Timko *et al*., 2012; Westwood 2013; Yoshida *et al*., 2016; Shahid *et al*., 2018; Clarke *et al*., 2019).

*Striga* is a genus of obligate, root parasitic plants within the Orobanchaceae (Parker & Riches 1993; Spallek *et al*., 2013). One species in particular, *Striga hermonthica* (Del.) Benth., is a notorious parasite of rain-fed rice, maize, sorghum and millets, leading to devastating losses in crop yields for resource-poor farmers in sub-Saharan Africa (SSA) (Scholes & Press 2008; Rodenburg *et al*., 2016). Control of *S. hermonthica* is extremely difficult as the parasite is an obligate outbreeder, with high fecundity, wide dispersal and a persistent, long-lived seed bank (Parker & Riches 1993) leading to a large effective population size (Huang *et al*., 2012). Resistant crop varieties are a crucial component of successful control strategies (Scholes & Press 2008) yet, even for crop varieties considered highly resistant, genetic variation within parasite populations is such that a few individuals can overcome host resistance responses and form successful attachments (Gurney *et al*., 2006; Cissoko *et al*., 2011). To develop crop varieties with durable resistance against *S. hermonthica*, it is vital to understanding fully, the repertoire, mode of action and genetic variability of parasite VFs that suppress or circumvent host defences (Timko et al., 2012; Rodenburg et al., 2017). Given the highly polymorphic populations of *S. hermonthica* and genetic diversity of the seed bank, we hypothesised that *S. hermonthica* is likely to possess suites of VFs that allow it to overcome layers of resistance in multiple host plant varieties. The aim of this study was to identify candidate genes encoding polymorphic VFs in *S. hermonthica*.

To achieve our aims we combined two complementary approaches. First, we assembled and annotated the genome of *S. hermonthica*, and developed a pipeline for computational prediction of putative secreted proteins (the secretome) and candidate VFs. The assembled genome was then used as a reference for an experimental, population genomics analysis, to compare DNA sequence variants in bulked (pooled) samples of *S. hermonthica* grown on a susceptible (NERICA-7) or resistant (NERICA-17) rice host (Fig. 1a i-ii). This allowed us to scan for loci in the *S. hermonthica* genome where the selection imposed by the resistant host had elevated the frequency of alleles contributing to successful colonisation (termed ‘virulence’ alleles) (Fig. 1 b-d). A similar approach was used to identify candidate genomic regions associated with resistance in *Solanum vernei* to the potato cyst nematode, *Globodera pallida* (Eoche-Bosy *et al*., 2017). The intersection between genes encoding predicted VFs and genes with highly significant allele frequency differences in the genome scan of *S. hermonthica*, revealed a set of candidate virulence loci encoding proteins with many functions, including cell wall modification, protease, or protease inhibitor, oxidoreductase and putative receptor-like protein kinase activities. Our results show that diverse strategies are used by *S. hermonthica* to overcome different layers of host resistance and suggest a polygenic basis of virulence in this parasite.

**Figure. 1.**
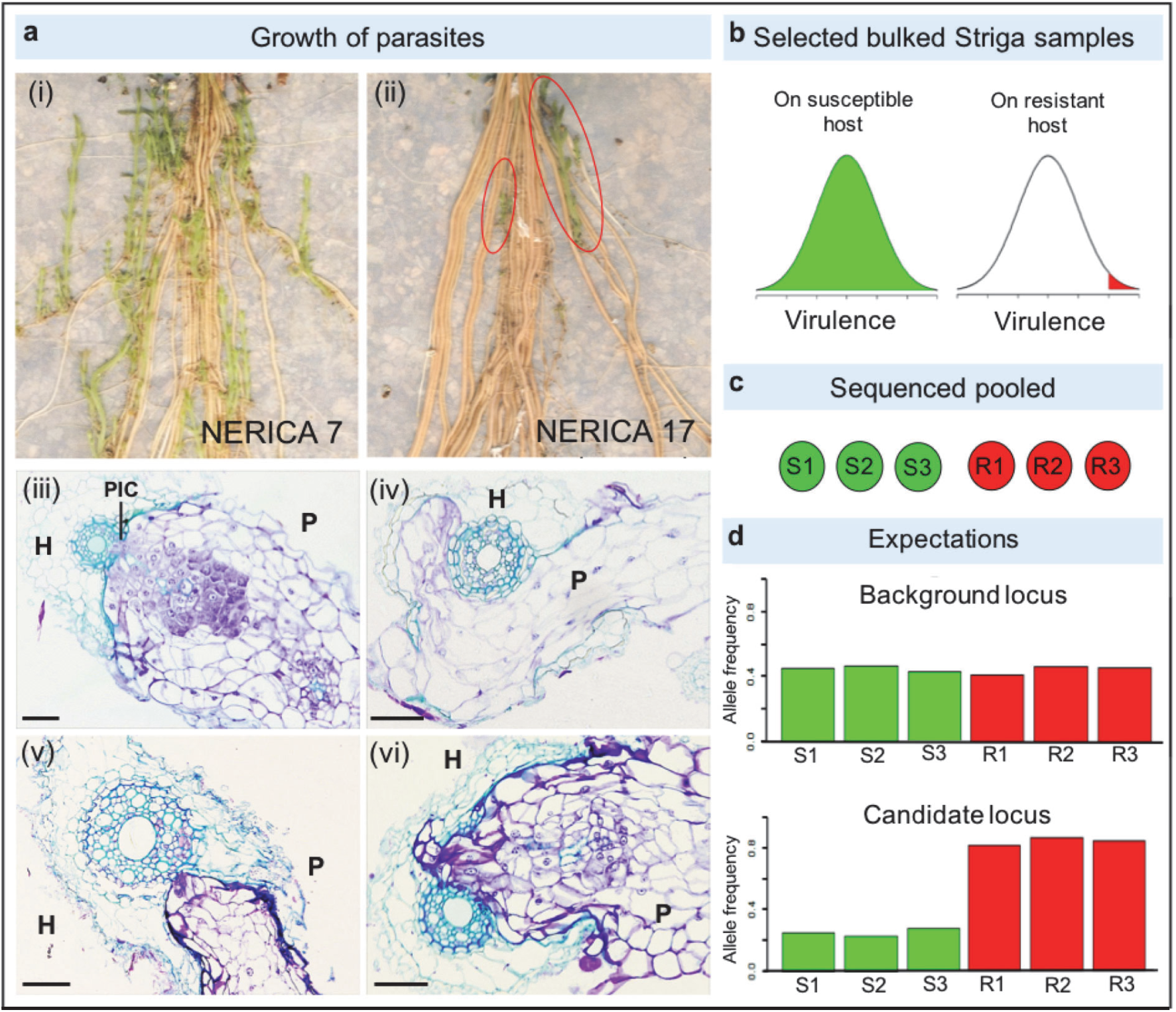
Experimental strategy for the identification of *Striga hermonthica* virulence loci. *Striga hermonthica* (Kibos accession) were grown on susceptible (NERICA 7) and resistant (NERICA 17) rice hosts (**a**). The whole rice root systems show many *S. hermonthica* individuals parasitising the roots of NERICA 7 (i) whilst only two individuals (red circles) were able to overcome the resistance response of NERICA 17 (ii) Scale = 1 cm. Transverse sections show *S. hermonthica* invading rice roots for a representative susceptible (iii) and resistant (iv–vi) interaction seven days post inoculation. In the successful host-parasite interaction parasite intrusive cells (PIC) have breached the endodermis and have made connections with the host’s xylem (iii). In the resistant rice variety several phenotypes are observed; The parasite invades the host root cortex but is unable to penetrate the suberized endodermis (iv, v); the parasite penetrates the endodermis but is unable to form connections with the host xylem (v). H = host root. P = parasite. Scale = 5 μm. Our experimental strategy was based on the prediction that many *S. hermonthica* genotypes would grow on NERICA 7 but only highly virulent genotypes would grow on NERICA 17 (**b**). Samples of 100 *S. hermonthica* plants were bulked to generate three sequencing pools from each host variety (**c**). We expected that background loci would not differ in allele frequency between pools, but virulence alleles (and neutral alleles in linkage disequilibrium) would have increased frequency in all pools from the resistant host, allowing us to identify candidate loci (**d**).

## Materials and Methods

### Collection and extraction of *S. hermonthica* DNA for genome and pooled sequencing

An accession (population sample) of *S. hermonthica* seeds was collected from individuals’ parasitising maize in farmers’ fields in the Kibos region of Kenya (0o 5’ 30.1272” S; 34 o 46’ 4.6416” E). To obtain *S. hermonthica* for genome sequencing and the bulked sample analysis (BSA), rice seedlings of the varieties, NERICA-7 and NERICA-17, were grown in rhizotrons and infected with germinated *S. hermonthica* seeds as described in (Gurney *et al*., 2006). Plants were grown in a controlled environment room with a 12 h photoperiod, a photon-flux density of 500 µmol.quanta.m^−2^.s^−1^ at plant height, a day / night temperature of 28 / 25 °C and 60 % relative humidity. For the construction of a reference genome, one *S. hermonthica* individual was randomly harvested from NERICA-7. For the pooled sequencing, 300 *S. hermonthica* individuals (> 30 mg in weight) were harvested from NERICA-7 and from NERICA-17, divided into 20 mg aliquots and immediately frozen in liquid nitrogen. The 300 individuals from NERICA-7 or NERICA-17 were divided into three pools of 100 individuals (three biological replicates). DNA was extracted from the six pools (see Methods S1) and samples were subjected to paired-end sequencing using an Illumina HiSeq machine at the Beijing Genomics Institute (BGI), China. The libraries, insert sizes and sequencing depth are shown in Table S1. DNA from the individual harvested from NERICA-7 for the production of a reference genome was sequenced on an Illumina HiSeq2500 sequencer at Edinburgh Genomics, UK. Six paired-end DNA libraries were constructed with different insert sizes (Table S1).

### De novo assembly of the *S. hermonthica* genome

Reads were cleaned and filtered as described in Methods S1. After filtering, ∼2.7 billion reads were generated from the short insert libraries and 0.76 billion reads from mate-pair libraries. This corresponded to ∼230 X and ∼54 X coverage of the *S. hermonthica* genome, respectively. The cleaned and filtered reads were used to assess the *S. hermonthica* genome size, repetitiveness and heterozygosity, compared with 12 other plant species (Table S2), in the module preQC, implemented in the software sga (https://github.com/jts/sga). This analysis showed *S. hermonthica* was highly heterozygous and therefore the software Platanus, which is specifically designed for highly heterozygous genomes, was chosen to assemble the *S. hermonthica* genome (Kajitani *et al*., 2014) (Table S3).

To further improve the *S. hermonthica* genome assembly, Chicago and Dovetail Hi-C libraries were prepared and sequenced at Dovetail Genomics, California, USA (https://dovetailgenomics.com/plant-animal/) (Table S3). For construction of Chicago libraries, DNA from the same *S. hermonthica* individual (used for initial sequencing) was sequenced on an Illumina HiSeq 2500 platform. For the Hi-C libraries, plant tissues from an F1 individual from a cross between the sequenced individual and another *S. hermonthica* individual (Kibos accession) were used for the library construction and sequencing. Sequences from both the Chicago and Hi-C libraries were used only to improve the contiguity of the initial genome assembly using the Dovetail HiRise Assembler software. RepeatModeler was used to generate a *S. hermonthica*-specific repeat library and RepeatMasker was then used to classify repeat elements in the genome. A repeat-masked version of the genome was used for annotation (Smit *et al*., 2008; 2013).

### Annotation of the *S. hermonthica* genome

The genome was annotated using three methods. Firstly, gene structures were inferred using a *S. hermonthica* transcriptome dataset of cDNAs collected from *S. hermonthica* individuals at eight developmental stages, generated by the Parasitic Plant Genome Project (PPGP) (Westwood *et al*., 2012; Yang *et al*., 2015). The reads were mapped onto the *S. hermonthica* genome assembly using TopHat to identify exon regions and splice positions (Trapnell *et al*., 2009). Transcriptome-based gene structures were predicted using Cufflinks (Trapnell *et al*., 2012) and candidate coding regions were then constructed in Transdecoder (https://github.com/TransDecoder/). Secondly, protein sequences from *Arabidopsis thaliana* (TAIR10), *Mimulus guttatus* (v2.0), *Solanum lycopersicum* (ITAG2.4), *Oryza sativa* (IRGSP1.0) and *Sorghum bicolor* (79), were used to determine consensus gene models in the genome. The protein sequences were mapped onto the *S. hermonthica* genome using TBLASTN and pairwise alignments were then input into Genewise (Birney 2004) to predict gene models in *S. hermonthica*. Thirdly, an *ab initio* method was used for *de novo* prediction of genes in the *S. hermonthica* genome using the software, Braker (Hoff *et al*., 2016). Finally, Evidence Gene Modeler was used to integrate various gene models from the transcript data, mapped proteins, and the predicted gene models from the *ab initio* method (Haas *et al*., 2008). The completeness of the gene set was assessed using BUSCO v5 using the 2,326 core orthologs from eudicots_odb10, with default settings.

### Functional annotation of the *S. hermonthica* proteome

Putative protein functions were assigned to *S. hermonthica* proteins using BLASTp analyses against the SwissProt and TrEMBL databases, and against the proteomes of *Arabidopsis thaliana* (version 30) and *Oryza sativa* (version 7). A BLASTp analysis was also conducted against the pathogen-host interaction database (PHI-base, version 4.2) (http://www.phi-base.org/index.jsp). BLASTp analyses were run locally using the NCBI BLAST package (version: ncbi-2.3.0+) and a hit was taken to be significant if e-value < 10^−5^, bit score and percentage identity > 30. Protein motifs and domains were determined by searching databases including Pfam, PATHER, GENE3D, CDD, PRINTS, PROSITE, ProDom and SMART with InterProScan Gene Ontology (GO) terms for individual proteins retrieved from the corresponding InterPro descriptions.

### Inference of orthogroups (OG)

Orthologous gene groups (OGs) were inferred using the software OrthoFinder v2 (Emms & Kelly, 2015). The number of genes per species for each OG was transformed into a matrix of Z-scores to quantify gene family expansion / contraction. The significance of expansion or contraction was determined using CAFE v4.2 (Han *et al*., 2013). Functional annotation of OGs was predicted based on sequence similarity to the InterPro protein family database. See full details in Methods S1.

### Prediction, analysis and refinement of the *S. hermonthica* secretome

Secreted *S. hermonthica* proteins were predicted using SignalP v 3.0 and 4.1 (Bendtsen *et al*., 2004; Petersen *et al*., 2011) (Fig. S1). Transmembrane spanning regions were identified using TMHMM2.0 (Krogh *et al*., 2001). Proteins with a secretion signal but without a predicted transmembrane helix were retained as the ‘secretome’. Pfam domains enriched in the *S. hermonthica* secretome compared with the rest of the proteome (non-secretome) were significant when the corrected p value was < 0.1, according to a Chi-squared test with a false discovery rate (FDR) correction for multiple testing (Benjamini *et al*., 1995). The initial secretome was then refined into subsets based on a series of structural and functional characteristics (Fig. S1) See Methods S1.

### Identification and analysis of candidate virulence loci using pooled sequencing data

The raw sequence reads from the six pools were trimmed and filtered for coverage (see Methods S1). The likelihood of the observed read counts for the two most common alleles, across the six pools was calculated according to equation 3 from Gompert and Buerkle (2011) to allow for the two levels of sampling associated with pooled sequencing data (sampling of reads and of individuals). We compared three allele-frequency models for each SNP using the Akaike information criterion (AIC): a null model with a single allele frequency for all pools, a control-virulent model with one frequency for the control pools (from the NERICA-7 host) and one for the virulent pools (from the NERICA-17 host) and a replicate model with a different allele frequency for each of the three pairs of pools (one control and one virulent) that were sequenced together. The control-virulent model was the model of interest while the replicate model was intended to check for consistency across pairs of pools. Therefore, two ΔAIC values were obtained: ΔAICcv = AICnull - AICcontrol-virulent and ΔAICrep = AICcontrol-virulent – AICreplicate. High positive values of ΔAICcv represent better fits than the null model and indicate significant differences between control and virulent pool types. SNPs with positive ΔAICrep values were likely to be affected by artefacts caused by sequencing methods and were excluded from the following analyses. All analysis steps were repeated independently for SNPs based on BWA and NOVOALIGN mapping as recommended by Kofler *et al*., (2016).

The effective population size in Striga is likely to be large (Parker & Riches 1993) and this is consistent with high diversity in our samples (overall mean π = 0.011). Therefore, we also expected that linkage disequilibrium would break down quickly. To define a suitable window size to search for regions potentially implicated in virulence, the extent of linkage disequilibrium in *S. hermonthica* was investigated (see Methods S1 for details). On the basis of this analysis, 1 kbp windows were used to detect genomic regions potentially associated with virulence on the basis of allele frequency differences between pools from the susceptible and resistant hosts.

Regions starting from 5kbp upstream of the start codon and ending no further than 2 kbp downstream of the stop codon of a gene were divided into 1 kbp-windows and the mean ΔAICcv across all the SNPs in each window was calculated. A permutation test was performed to obtain the probability of observing the mean ΔAICcv value, or higher, for each window based on the distribution of ΔAICcv across the regions as a whole (see Methods S1 for details). Finally, we retained genic regions (defined as regions from 2 kbp upstream of the start codon to the 1 kbp window containing the stop codon) for which this probability was less than or equal to 2×10^−5^ for both the BWA and NOVOALIGN analyses in any window. This cut-off was chosen to provide experiment-wide significance given the number of protein-coding genes in the analysis (29,518). In the secretome, a more relaxed cut-off of 1×10^−4^ was used to reflect the prior expectation that the secretome would be enriched with pathogenicity-related genes and the smaller number of genes in this set (3,375). Thirty-two genes met this criterion for both Novoalign and BWA (Data S1). In addition, six genes encoding putative secreted proteins that passed the 1×10^−4^ cut-off for either Novoalign or BWA were included in the candidate set because they either contained large numbers of non-synonymous SNPs or contained high impact SNPs that can alter protein structure (e.g. due to protein truncation) (Data S1).

Two population statistics were calculated for each genic region in the control pool using the software Popoolation (Kofler et al., 2011). These were nucleotide diversity (π) and Tajima’s D, a statistic describing the allele frequency spectrum used for testing whether a DNA sequence is evolving under a process that departs from the standard neutral model, such as selection or demographic change (Tajima, 1989). See Methods S1 for details.

### Analyses of candidate virulence genes

The candidate virulence genes were categorised into functional groups based on the annotations of the closest matching homologs from the *A. thaliana* and *O. sativa* proteomes, as well as the Pfam domain annotations. For each gene, the numbers of SNPs were counted for the promoter region (within 2 kbp upstream of the start codon), the intronic region and coding region, and the numbers of non-synonymous SNPs were determined. To quantify the allele frequency differences between control and virulent pools for these candidate virulence genes, the proportion of SNPs with high fixation index (F_ST_) values in the significant window was calculated (see Methods S1).

### Expression profiling of candidate virulence genes

Expression profiles for candidate virulence genes were determined for *S. hermonthica* collected at 2, 4, or 7 days post infection from the roots of NERICA-7 rice plants (full details are provided in Methods S1). In addition, unattached *S. hermonthica* haustoria were induced *in vitro* by the addition of 10 μM DMBQ (Fernández-Aparicio *et al*., 2013). Cleaned reads were mapped to the *S. hermonthica* genome using Tophat2, version v2.0.12 and quantified with HTSeq (version 0.6.1). FPKM values for each gene at each time point were used to calculate a fold change in expression relative to the haustorial sample and significance assessed with a one-way ANOVA. For each gene, log2 fold expression values, across the time points, were centred around 0 and scaled by the standard deviation for plotting as a heatmap using the pheatmap function in R. Further details are provided in Methods S1.

## Results

### The *S. hermonthica* genome is very heterozygous

We obtained a single population of *S. hermonthica* seeds from farmer’s fields in Kibos, Kenya and infected a highly susceptible rice variety, NERICA-7 (Fig. 1a). The genetic diversity of the seed population is reflected in the subtle differences of flower colour and morphology of attached parasites (Fig. 2a). We sequenced, assembled and characterised the genome of a single individual from this population, which to our knowledge, represents the first genome assembly for *S. hermonthica*. The genome size was estimated by K-mer analysis to be 1,475 Mbp, (Fig. 2b). This agrees closely with a previous flow cytometry-based estimate (Estep *et al*., 2012) and is more than twice the size of the recently sequenced genome of *S. asiatica* (Yoshida *et al*., 2019). The assembly consisted of 34,907 scaffolds > 1 kbp in length, with an N50 of 10.0 Mbp and 29 scaffolds making up half of the genome size (Table S3). The *S. hermonthica* genome was remarkably heterozygous (overall mean π = 0.011) (Fig. 2 b,c) when compared with other parasitic and non-parasitic plant genomes, likely reflecting the fact that it is an obligate outbreeding species. In addition, the genome contained a large proportion (69%) of repetitive DNA (Fig 2 b,c), dominated by long terminal repeat (LTR) elements (Table S4), a pattern also found for the shoot-parasitic plants, *Cuscuta australis* and *C. campestris* (Sun *et al*., 2018; Vogel *et al*., 2018) and the closely related parasitic plant *S. asiatica* (Yoshida *et al*., 2019). As expected, the density of repetitive elements along each scaffold negatively correlated with the density of protein-coding genes (Fig 2c). In total, 29,518 protein-coding genes were predicted from the *S. hermonthica* genome, which was comparable to *S. asiatica* (34,577), the closely related non-parasitic plant *Mimulus guttatus* (28,140) and to *Arabidopsis thaliana* (27,416) (Table S5).

**Figure. 2.**
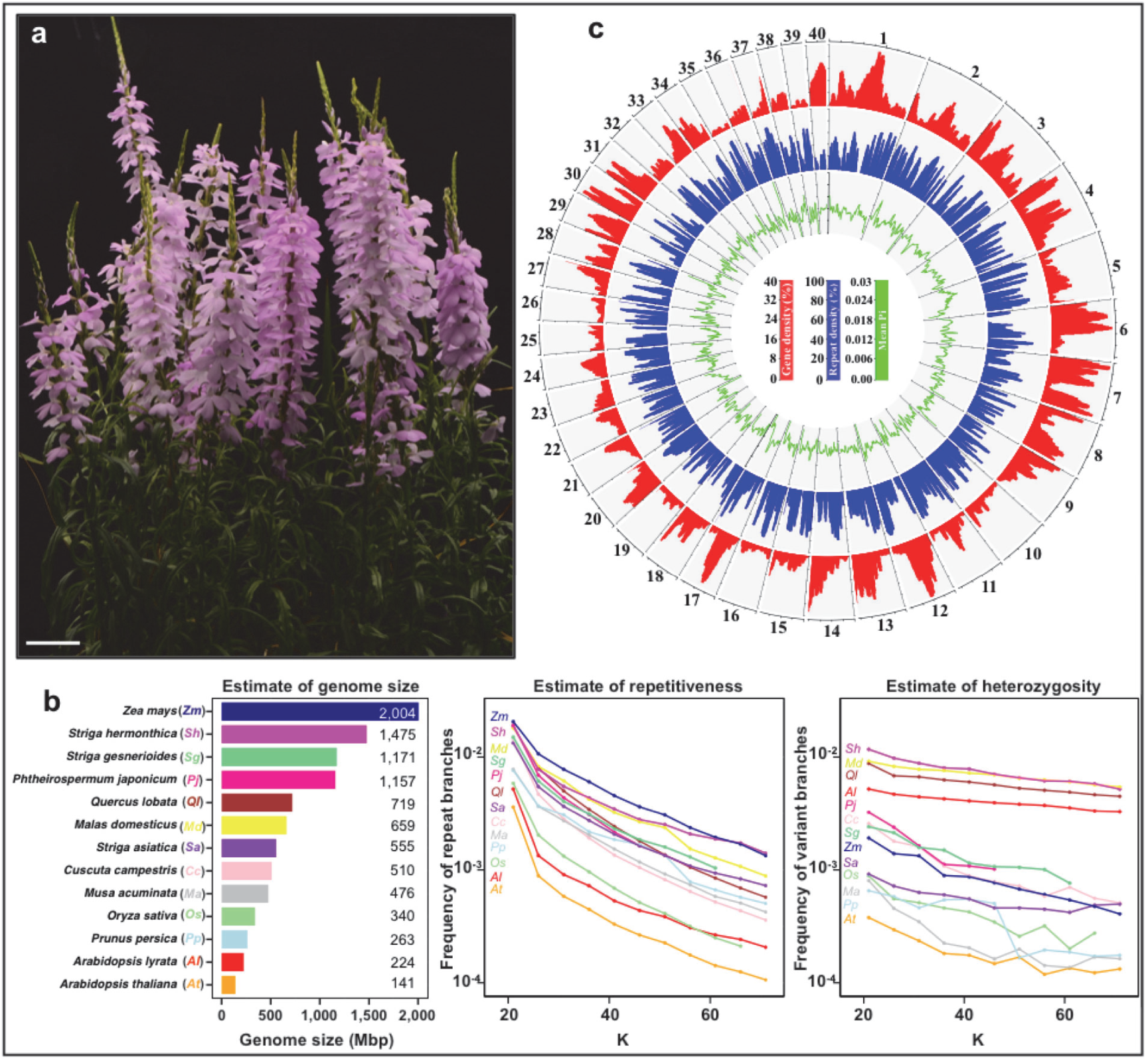
*Striga hermonthica* is an obligate outbreeding parasitic plant with a highly heterozygous and repetitive genome. **a**, Flowering *S. hermonthica* growing on the rice host, NERICA 7, derived from a seed batch collected from the Kibos region of Kenya. Scale = 5 cm. **b**, Comparison of genome size, heterozygosity and repetitiveness between *S. hermonthica* and 12 other plants. The estimate of the genome size (Mbp) was based on k-mer count statistics. The estimate of heterozygosity was based on variant branches in the k-de Bruijn graph. The repetitiveness of the genomes was based on frequency of repeat branches in the k-de Bruijn graph. K: k-mer length. **c**, Genomic features calculated in 1 Mbp windows with a slide of 250 kbp for the largest 40 scaffolds in the *S. hermonthica* genome assembly. Outer bar plot (red): gene density (percentage of the window comprised of genic regions). Mid bar plot (blue): repeat density (percentage of window comprised of repetitive sequence). Inner line plot (green): nucleotide diversity (mean Pi for genic regions). Axes tick marks around plot circumference denote 4 Mbp. Vertical axis tick marks are defined in the centre.

BUSCO analysis of gene set completeness (Waterhouse *et al*., 2018), showed 87.3% of 2,326 conserved single-copy orthologs in eudiocotyledons were complete in the *S. hermonthica* genome, similar to that found in *S. asiatica* (88.7%) (Fig. 3; Table S6). Of the BUSCOs not found in the *S. hermonthica* genome, over half were also absent from the *S. asiatica* genome (Table S6). Both *Striga* spp. share missing BUSCOs that are present in the genome of the closely related non-parasitic *Mimulus guttatus* (Fig. 3b; Table S6). Similarly, two shoot holoparasites, *C. australis* and *C. campestris*, with a BUSCO completeness of 81.0 and 81.7% respectively, also shared many missing BUSCOs that were present in the genome of their non-parasitic relative, *Ipomea nil* (Fig. 3c). This is consistent with previous findings suggesting some missing BUSCOs are likely to be a result of the parasitic lifestyle (Sun *et al*., 2018; Vogel *et al*., 2018; Yoshida *et al*., 2019; Cai *et al*., 2021).

**Figure. 3.**
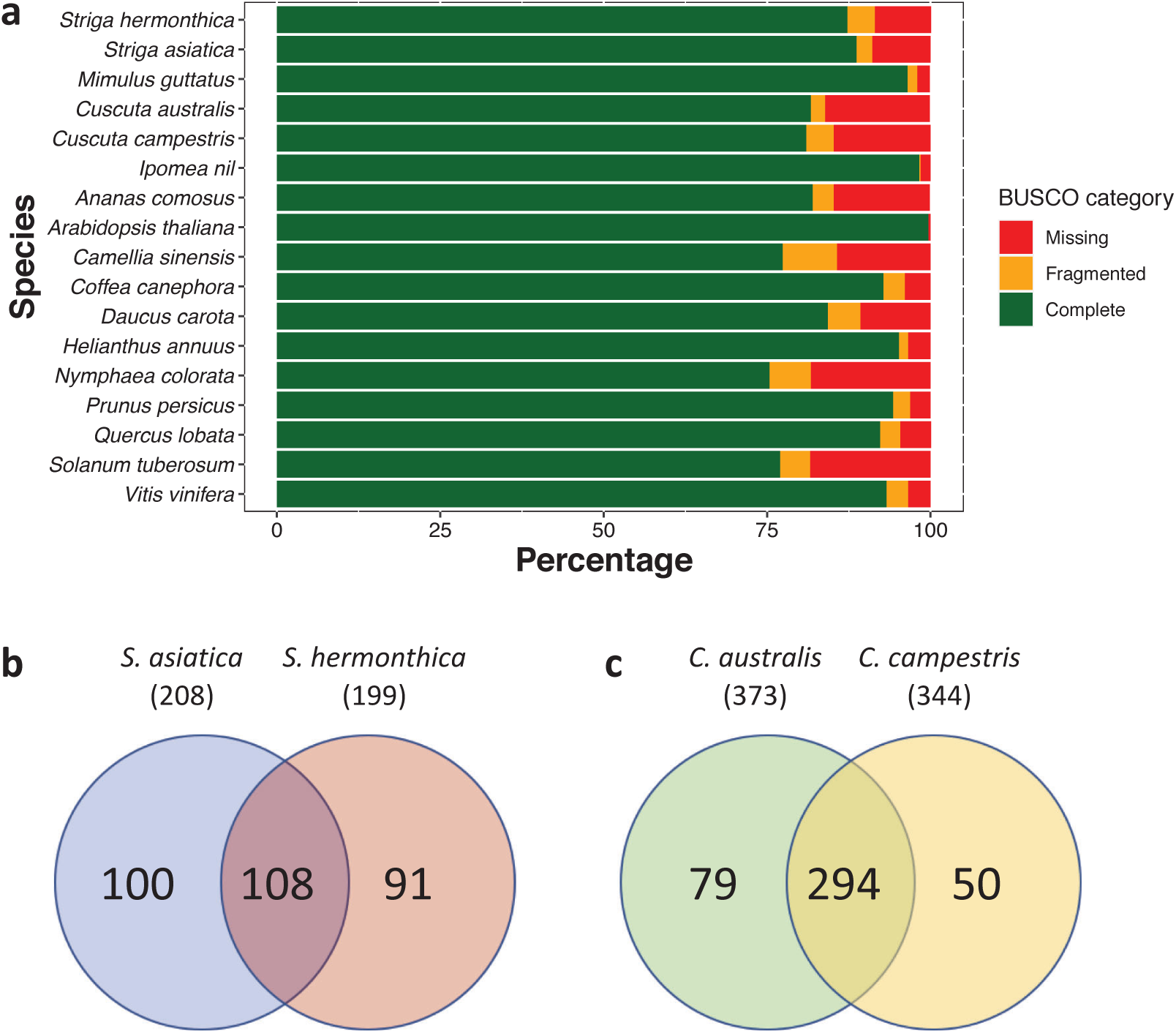
**a**, BUSCO completeness analysis for *Striga hermonthica* genome, compared with 16 other published plant genomes. The number of missing BUSCOs for two Striga **b** and two Cuscuta species **c**. The overlap shows genes that are missing from both Striga or Cuscuta species respectively.

Comparative analysis of orthologous gene groups (orthogroups) between *S. hermonthica* and 12 other plant species identified 22,624 orthogroups in total, of which 12,278 contained *S. hermonthica* genes. Of these, 327 were significantly expanded and 104 were contracted in the *S. hermonthica* genome (Fig. 4a). Expanded orthogroups included the *α*/*β*-hydrolase family, recently shown to have undergone duplication in *S. hermonthica* (Toh *et al*., 2015), as well as numerous F-box, leucine-rich repeat and protein kinase domain-containing proteins (Fig. 4b). Of particular interest in the context of pathogenicity were *S. hermonthica*-specific orthogroups annotated as papain family cysteine proteases, xylanase inhibitors and trypsin and protease inhibitors (Fig. 4b). Both proteases and protease inhibitors function in a wide range of plant-plant parasite interactions and may act offensively, by degrading host proteins, or defensively, by inhibiting host defence enzymes (Bleischwitz *et al*., 2010; Mueller *et al*., 2013).

**Figure. 4.**
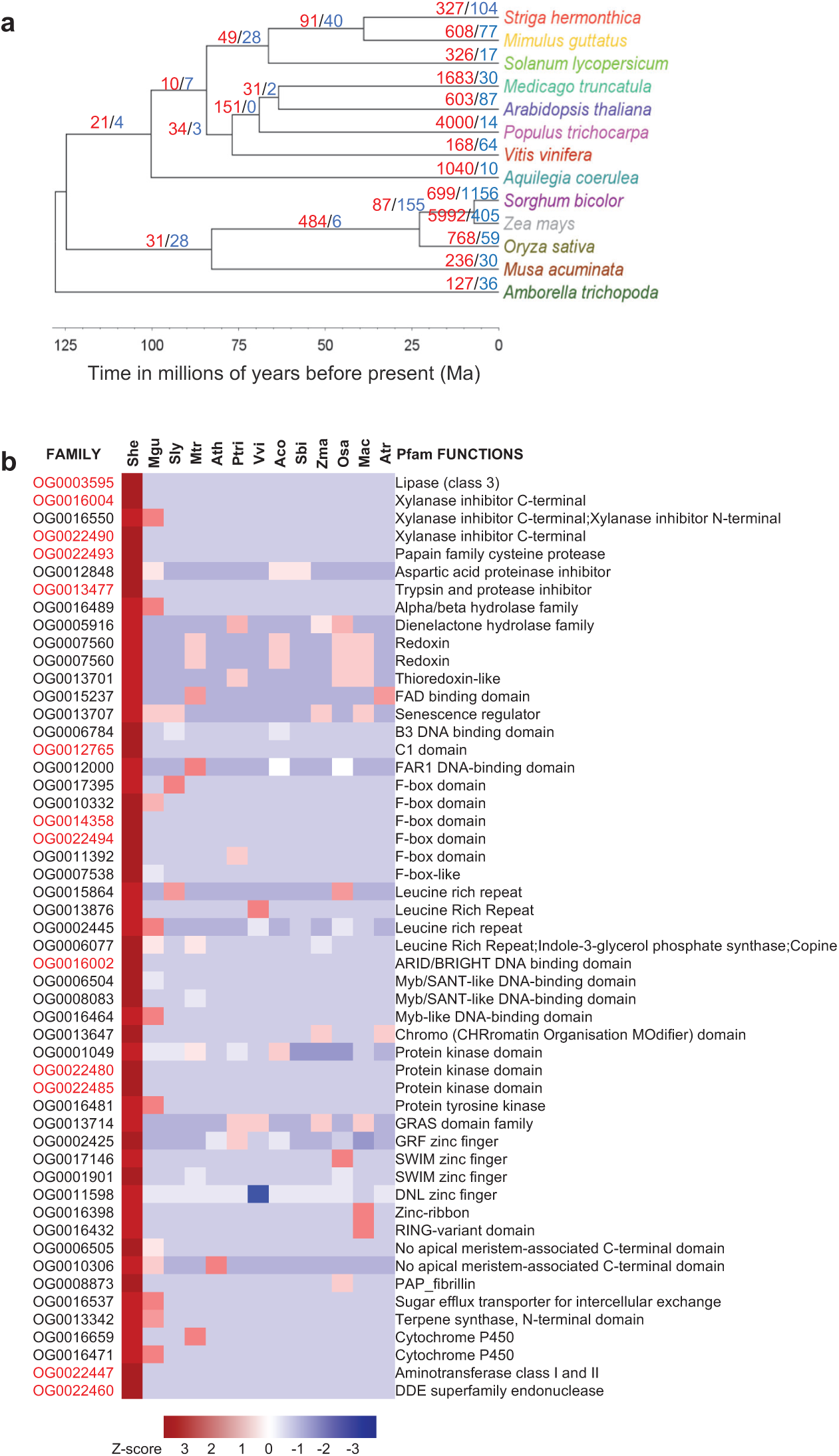
Orthogroup analyses. **a** A time tree for *S. hermonthica* and 12 other species generated in MEGA, based on 42 single-copy genes inferred from OrthoFinder. The number of significantly expanded (red) and contracted (blue) orthogroups based on CAFE analysis are shown above the branches. **b** Significantly expanded orthogroups in *S. hermonthica*, after removing proteins encoded as transposable elements, compared to 12 other plant species. Orthogroups only found in *S. hermonthica*, have family names in red. Higher Z-scores indicate the orthogroups are more expanded in a species while lower Z-scores indicate the orthogroups are more contracted in a species.

### The *S. hermonthica* secretome

One way that parasite proteins can interact with host biology is through parasite-directed secretion. We identified 3,375 putatively-secreted proteins in *S. hermonthica* (11.4 % of the proteome) (Fig. S1), many of which were homologous to *A. thaliana* secreted proteins (Table S7), providing experimental evidence for secretion into the extracellular space. On average, the *S. hermonthica* secreted proteins were both significantly smaller and had a higher percentage of cysteine residues compared with the rest of the proteome (Fig. 5 a, b). Genes encoding secreted proteins tended to be more clustered (within 15 kbp of their nearest neighbour) compared to all genes in the genome (p < 10^−4^, 10^5^ permutations) (Fig. S2) suggesting they are likely to be arrayed in tandem and belong to large gene families (Elizondo *et al*., 2009). Functionally, the secretome was rich in protein domains involved in cell wall modification (e.g. endoglucanases, cellulases, pectinesterases, expansins, and pectate lyases), protease activity (e.g. papain-like cysteine proteases, aspartic proteases, and subtilase proteases) and oxidoreductase activity (peroxidases, copper oxidases, and cytochrome p450 proteins) (Fig. 5c, Figs. S3 and S4). The cytochrome P450 domain, for example, was particularly frequent in the *S. hermonthica* secretome (3.13% of protein domains) compared with the rest of the proteome (0.25% of protein domains) (Fig. S3). Three other highly-abundant protein domains in the secretome were described as copper oxidases (Fig. S3) and are commonly found in laccases that are involved in the generation or breakdown of phenolic components, such as lignin (Kwiatos *et al*., 2015). Small cysteine-rich proteins are common characteristics of VFs from a range of phytoparasites (Saunders *et al*., 2012; Lu *et al*., 2016). In *S. hermonthica*, 183 such proteins were identified (Fig. 5a) and were similar to proteins annotated as carbohydrate binding X8 domain-containing proteins, protease inhibitor/lipid transfer proteins, PAR1-like proteins, pectinesterases, RALF-like proteins and thaumatin-like proteins (Fig. S4), many of which are likely to play a role in host-Striga interactions (Yang *et al*., 2015; Yoshida *et al*., 2019).

**Figure. 5.**
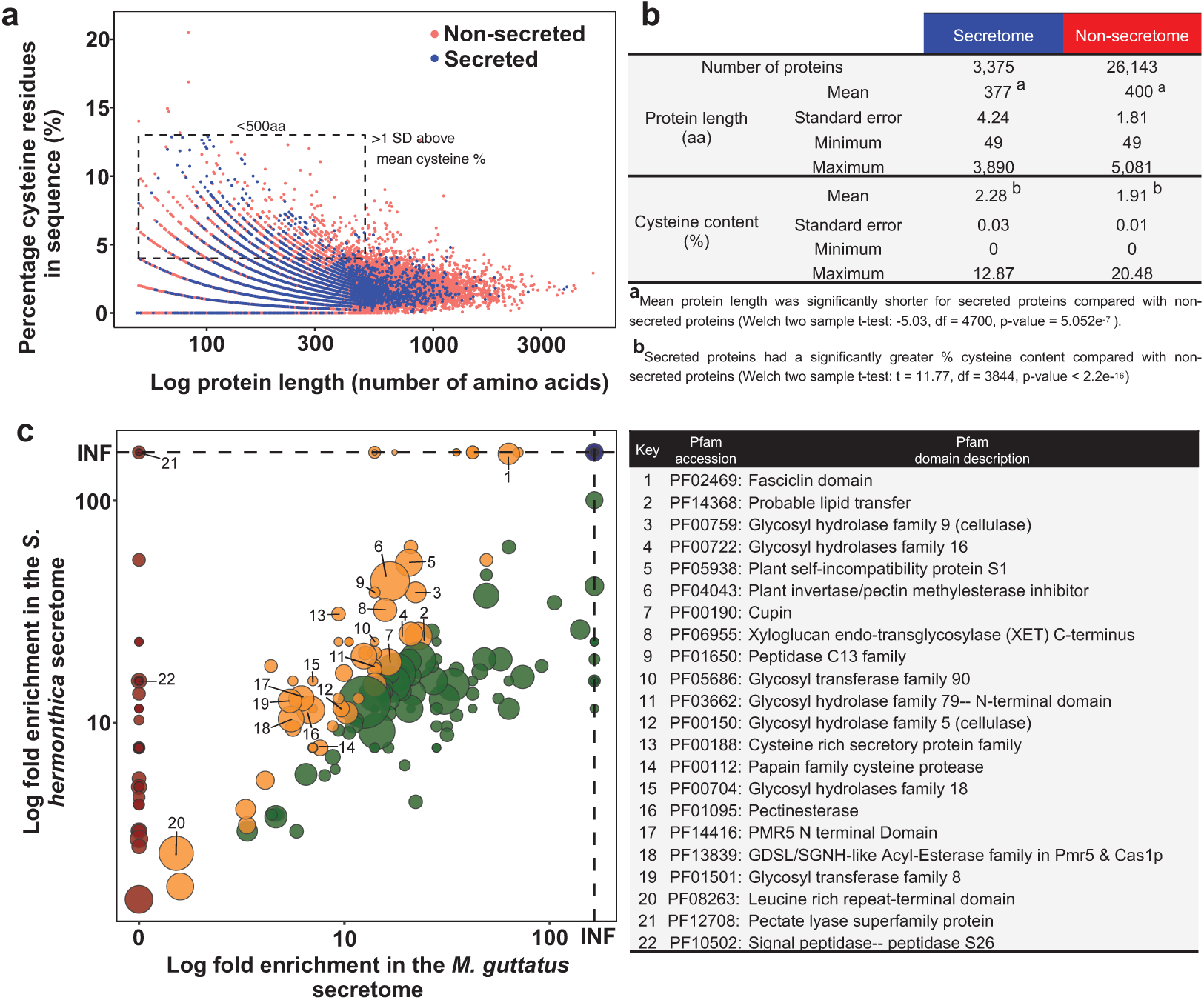
*Striga hermonthica* secretome. **a**, Relationship between protein length (log scale) and cysteine content (as a % of total amino acid number) for putatively-secreted (blue) and non-secreted (red) proteins in the *S. hermonthica* proteome. Secreted proteins < 500 amino acids in length and with a cysteine % > 1 standard deviation above the mean, were selected as a subset of small, cysteine rich proteins. **b**, Descriptive statistics for length and cysteine content for secreted and non-secreted proteins. **c**, Pfam domains enrichment (log fold-change) in the *S. hermonthica* secretome, relative to the proteome as a whole, compared to the corresponding enrichment in the *Mimulus guttatus* secretome. INF denotes infinite enrichment (Pfam domain only found in the secretome). Points above the 1:1 diagonal were enriched more in the *S. hermonthica* secretome relative to *M. guttatus* and have been coloured accordingly. Red symbol: domains only enriched in the *S. hermonthica* secretome. Yellow symbol: domains enriched more in the *S. hermonthica* secretome than in the *M. guttatus* secretome. Green symbol: domains enriched more in the *M. guttatus* secretome than in the *S. hermonthica* secretome. Blue symbol: domains present only in the secretome in both species. Sizes of the points were weighted according to the frequency of occurrence of each Pfam domain in the *S. hermonthica* secretome. Annotations for the most significantly enriched of the Pfam domains (p < 0.01) that were also enriched more in the *S. hermonthica* secretome relative to the *M. guttatus* secretome, are given in the accompanying table with their functional descriptions.

We identified several protein domains in the *S. hermonthica* secretome that were enriched to a higher degree than observed in the secretome of the closely-related non-parasitic plant, *M. guttatus* (Fig. 5c, Fig S3, Data S2), suggesting these functions are relevant to the parasitic lifestyle. The xyloglucan endotransglycosylase (PF06955) domain, for example, was found in 17 *S. hermonthica* proteins (Fig. 5c, Fig. S4).

Xyloglucan endotransglucosylases / hydrolases (XETs) have the potential to modify either the parasite or host cell walls (or both) during parasitism (Olsen & Krause 2017). XETs are secreted from the haustoria of the parasitic plant *Cuscuta reflexa* during a susceptible interaction on its host *Pelargonium zonale*, contributing towards pathogenicity (Olsen & Krause 2017). Pectate lyase superfamily (PF12708) and pectinesterase (PF01095) domains were enriched in the secretome of *S. hermonthica* compared to *M. guttatus* and may act as VFs to modify host, or parasite, pectin during penetration. We found a battery of different carbohydrate-active glycosyl hydrolase (GH) domains that were enriched in the *S. hermonthica* secretome (Fig. 5c, Fig. S3). Eight *S. hermonthica* proteins were annotated as cellulases of the GH5 family (containing domain PF00150) (Fig. S4) and were similar to secreted cellulases that function as VFs in some phytoparasitic nematodes (Smant *et al*., 1998). The degradation of cellulosic β-1,4-glucans has been observed in susceptible sorghum roots infected by *S. hermonthica* (Olivier *et al*., 1991) and may be mediated by these secreted enzymes to facilitate the migration of *S. hermonthica* intrusive cells between host root cortical cells. The identification of many putatively secreted VFs in the *S. hermonthica* genome, that are likely to modify host plant cell walls, raises an interesting question about how such proteins are targeted to avoid damaging the parasite’s own cell walls.

### Population genomic analysis to identify candidate virulence loci

Our experimental system allowed us to identify a subset of VFs with genetic variation relevant to the ability to infect some host genotypes and not others. Hundreds of *S. hermonthica* individuals were harvested from either a very resistant (NERICA-17) or susceptible (NERICA-7) rice cultivar, and pools of these individuals were subjected to genome resequencing. After aligning the reads to our reference genome, we detected 1.8 million SNPs in genic regions. These genic regions were split into 150,741 1 kbp windows and of these, 194 (0.13%) had extreme and consistent allele frequency differences between the bulked pools of *S. hermonthica* selected on the resistant *versus* the susceptible hosts (Fig. S5; Data S1). These highly differentiated windows were located in 190 genes. These candidate loci potentially encode virulence factors with allelic variants, influencing either structure or expression that contribute to the ability of some individuals to parasitise NERICA-17. As expected for an outbred parasite with a large population that encounters multiple host species and genotypes, many loci were detected and they cover a range of predicted functions. Of these candidate VFs, 152 were not predicted to be secreted and were assigned to a wide range of functional categories, including putative transcription factors, hormone signalling pathways, transporters, repeat-containing proteins and a number of proteins of unknown function (Fig. 6a; Data S1). Some of these proteins may function to protect the parasite against host defences and facilitate growth on the resistant rice variety. In addition, some may enter the host by non-traditional pathways, for example, via the host-parasite xylem connections. One sixth (24) of these non-secreted proteins had sequence similarity to proteins in the Pathogen-Host Interaction database (Winnenburg *et al*., 2007). These included *S. hermonthica* proteins with sequence similarity to a putative leucine-rich repeat protein from *Ralstonia solanacearum*, a mitogen-activated protein kinase from *Ustilago maydis*, a calreticulin-like protein from *Magnaporthe oryza* and a cytochrome P450 from *Bursaphelenchus xylophilus* (Data S1).

**Figure. 6.**
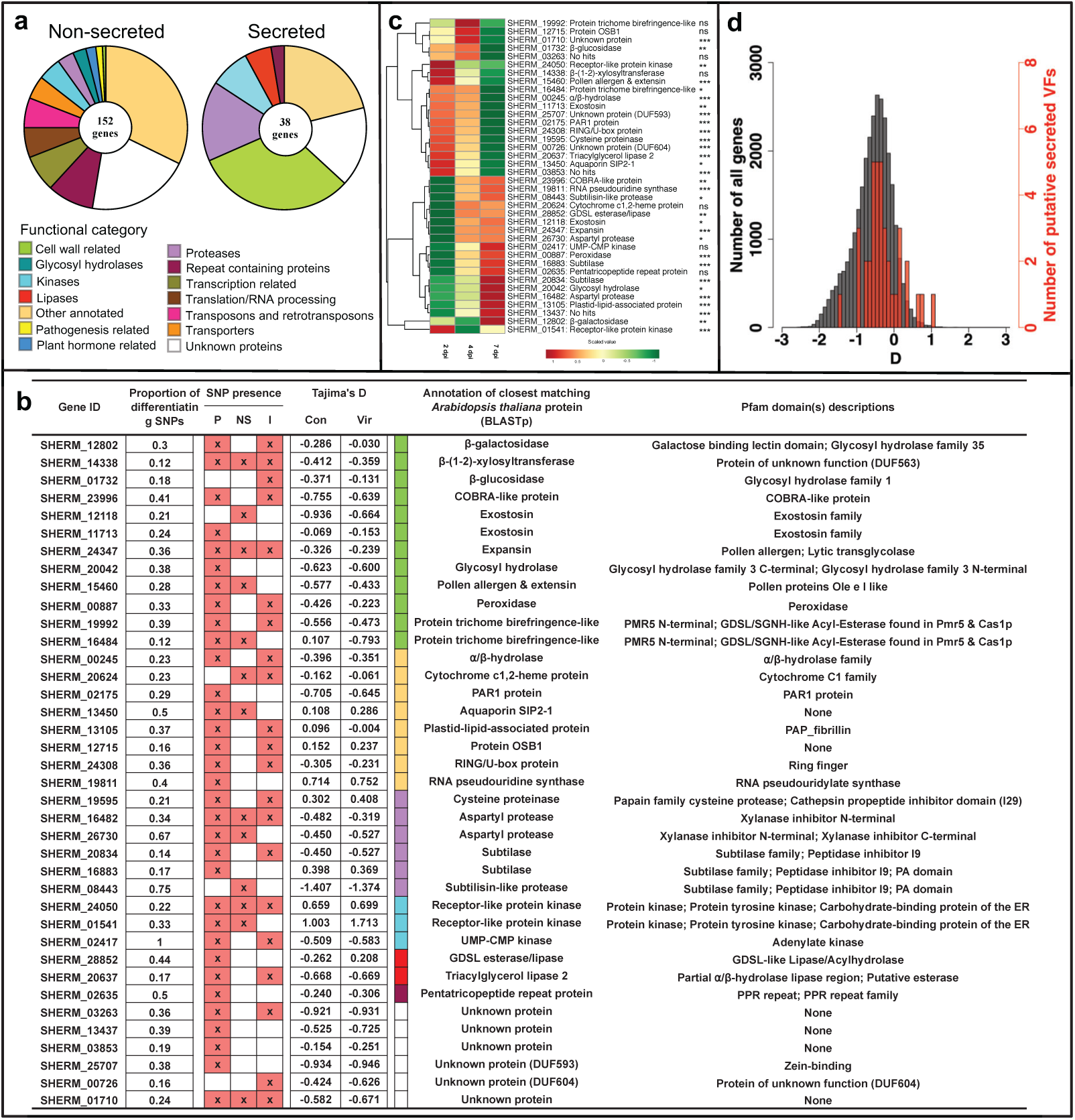
Identification of *Striga hermonthica* genes that display significant allele frequency differences between pools of individuals parasitising the susceptible rice variety (NERICA 7) and those that successfully parasitise the resistant rice variety (NERICA 17). **a** Functional categorisation of non-secreted proteins and secreted, candidate virulence factors (VFs). **b** The 38 genes encoding putative secreted *S. hermonthica* proteins with their associated measure of differentiation (proportion of differentiating SNPs within the significant window) between the control and virulent sets of pools. The presence of SNPs in the promoter region (P), non-synonymous SNPs in the coding region (NS) and those in the intronic regions (I) are indicated with an X. The annotation of the closest matching *Arabidopsis thaliana* protein is shown along with coloured boxes that correspond to the functional category assigned in the pie chart in **a**. Tajima’s D was calculated for individuals grown on NERICA 7 (Con) or NERICA 17 (Vir). **c**. Clustered gene expression profiles of the 38 candidate VFs in *S. hermonthica* haustoria parasitising NERICA 7 at 2, 4 and 7 days post-inoculation (dpi). Log_2_ fold change in expression is shown relative to expression levels in haustoria induced *in vitro*. The gene IDs and putative functions based on best BLASTp hit against the *A. thaliana* proteome correspond with part **b**. Significant changes in gene expression in haustoria during the infection time course are shown *** (p < 0.001); ** (p < 0.01); * (p < 0.05); ns non-significant (ANOVA). **d**. Comparison of Tajima’s D for the 38 putative VFs (red) and all the genes in the genome (grey) for the control pools.

The remaining 38 VFs were members of the *S. hermonthica* secretome and represent particularly strong candidates associated with the ability to parasitise NERICA-17 successfully (Fig. 6a,b, Data S1). These genes were categorised into six functional groups, the largest of which contained 12 genes associated with cell wall modification (Fig. 6a,b), including genes encoding an expansin protein, a COBRA-like protein, a β-(1-2)-xylosyltransferase, two trichome birefringence-like (TBL) proteins, a pollen Ole e allergen and two exostosin family proteins, all of which can function to modify the extensibility or other mechanical properties of plant cell walls (Li 2003; Qin *et al*., 2004; Honaas *et al*., 2013; Mitsumasu *et al*., 2015) (Fig. 6b). Groups of genes annotated as proteases (6 genes including subtilases, aspartyl proteases, and a cysteine proteinase), lipases (3 genes) and kinases (3 genes) were also found. The proteases were always associated with an inhibitor protein domain (Fig. 6b). For example, the putative aspartyl proteases possessed one or more xylanase inhibitor domain(s) (Fig. 6b). There were also eight genes encoding proteins with a range of putative functions, including a PAR1-like protein, a probable aquaporin, an *α*/*β*-hydrolase and two receptor-like protein kinases (Fig. 6b). In addition, a further six genes were annotated as proteins of unknown function (Fig. 6b).

The 38 candidate VFs were investigated in more detail by quantifying changes in gene expression in haustoria at critical stages of parasite development on the susceptible rice variety NERICA-7 by inspecting the distribution of SNPs throughout the promoter and genic regions, and testing for signatures of historical selection. Gene expression was measured in an independent experiment (Fig. 6c). Changes in gene expression of attached haustoria were measured relative to gene expression in haustoria generated *in vitro*. At 2 days after inoculation of the host root, parasite haustoria were attached and parasite intrusive cells had penetrated into the host root cortex. By day 4, the parasite intrusive cells had penetrated between the endodermal cells and by day 7 had formed connections with the xylem vessels of the host, providing direct access to host resources (Fig. 1a iii).

Prior to attachment to the host, some of the genes encoding candidate VFs were not expressed in haustoria (e.g. subtilase gene (SHERM_16883) and subtilisin-like protease (SHERM_08443) or were expressed at very low levels (e.g. the peroxidase (SHERM_00887), glycosyl hydrolase (SHERM_(20042), both aspartyl proteases (SHERM_16482 and SHERM_26730) and an unknown protein (SHERM_03853) (S3 Data). However, all 38 genes were expressed in haustoria during the early stages of infection of the susceptible host, NERICA-7 (Fig. 6c; Data S3). There were two main patterns of gene expression. Firstly, 21 genes, including those mentioned above, had low levels of expression in haustoria 2 days post infection, followed by an increase in expression as infection progressed (Fig. 6c; Data S3). In contrast, 17 genes were highly expressed in haustoria 2 days post infection and expression then decreased progressively with time, e.g. genes encoding β-glucosidase, β-(1-2)-xylosyltransferase, and TBL protein SHERM_06484, all of which modify cell walls. The cysteine protease, PAR1, ɑ/β-hydrolase and aquaporin genes also exhibited a similar expression profile (Fig. 6c; Data S3).

Most of the 38 genes had significantly differentiating SNPs in their promoter regions (from the start site to 2 kbp up-stream). Some of these SNPs may lead to a change in the regulation of gene expression (Fig 6b). Some genes, for example, the gene encoding the pollen Ole e allergen protein (SHERM_15460), one of the exostosin family proteins SHERM_12118), a probable aquaporin SIP2-1 (SHERM_13450) and one of the two protein TBL genes (SHERM_16484), also had non-synonymous SNPs in the coding region (Fig. 6b) that may result in functional differences between the alleles of these genes in individuals infecting NERICA-7 and NERICA-17. Finally, SNPs were also found within predicted intron regions in many of the genes (Fig. 6b).

The co-evolutionary interactions between hosts and parasites can generate balancing selection (Frank 1993). We predicted that genes contributing to virulence would tend to have a history of balancing selection because of the diverse range of hosts used by *S. hermonthica*. To test this prediction, we compared Tajima’s D between candidate loci and the rest of the genome, expecting to see more positive values (Charlesworth 2006). We used the pools from the susceptible host for this comparison because they represented the Striga population as a whole. As predicted, the 152 candidate loci in the *S. hermonthica* proteome (Fig S6) and the 38 candidate loci in the secretome (Fig. 6d) had significantly elevated Tajima’s D, on average, compared to all the genes in the genome (p < 0.0001 and *p* < 0.0003, respectively; 10^5^ permutations). Some loci had particularly high Tajima’s D values, for example the two receptor-like protein kinases (Fig. 6b). Interestingly, some loci showed large differences in Tajima’s D between the control and virulent *S. hermonthica* pools with the largest difference seen for the TBL gene (SHERM_16484) with a negative ΔD (D_Vir_ - D_Con_) of -0.9. This suggests strong selection resulting in one common haplotype in the virulent pools in contrast to two or more haplotypes at intermediate frequencies in the control pools. There were also large positive ΔD values: 0.71, 0.16 and 0.20 for one of the putative receptor-like protein kinases SHERM_01541, one of the aspartyl proteases, SHERM_16482, and the peroxidase SHERM_00887, respectively. This suggests that a rare haplotype in the control pools is present at intermediate frequency in the virulent pools. Overall, these changes indicate that selection on the resistant host caused changes in frequency of multi-SNP haplotypes at these loci, haplotypes that may have been created by areas of low recombination or by recent invasion of new variants under positive selection (Cutter & Payseur 2013) and which underlie the ability of some *S. hermonthica* individuals to overcome resistance in NERICA-17.

## Discussion

Plants secrete proteins involved in many biological functions, from nutrient acquisition, to development and defence (Li 2003; Cook *et al*., 2015). However, unlike most plants, in parasitic plants such as *S. hermonthica* a subset of secreted proteins is likely to function as VFs and contribute towards parasite fitness by facilitating host colonization (Timko *et al*., 2012). We used a combination of *in silico* prediction of secreted proteins and pooled sequencing of parasites derived from susceptible and resistant rice hosts, both facilitated by the first available genome assembly, to identify a set of candidate VFs. These are secreted proteins encoded by genes that had extremely different allele frequencies between replicated pools derived from susceptible and resistant hosts, suggesting strong selection for particular variants that facilitate successful colonisation despite host resistance. This experimental approach has not been applied previously to investigate virulence of Striga, or any other parasitic plant. Its success here paves the way to application of similar methods to other host-parasite combinations, providing vital information on virulence mechanisms and their genetic variability within and between parasitic plant populations from different regions of Africa, and so underpinning the development of sustainable control strategies.

Our list of 38 candidate, secreted, VFs points to key functions involved in pathogenicity, including oxidoreductase, receptor-like protein kinase, protease and protease inhibitor, and cell wall modification activities. The latter is consistent with growing evidence that cell-wall modification is a critical step in plant invasions by many different parasites including parasitic plants. Recently, the structural integrity of lignin was shown to be a crucial component of resistance in roots of the rice variety Nipponbare to infection by *S. hermonthica* (Mutuku *et al*., 2019). In our study the host cell wall is clearly involved in resistance in NERICA-17. Most *S. hermonthica* individuals from the Kibos population were unable to penetrate the root endodermis or, if they breached the endodermis, they were unable to establish functional connections to the host xylem vessels (Fig. 1a iv-vi). Consistent with this, the largest category of our candidate, secreted VFs included a putative peroxidase, an expansin, pollen allergen-like proteins, a β-glucosidase, a β (1-2) xylosyltransferase, and a TBL protein, all of which function to modify cell walls. The TBL protein, SHERM_16484, had a strikingly different Tajima’s D in the control pool compared to the value in the virulent pool, consistent with selection favouring one haplotype on the resistant NERICA-17, out of several haplotypes present in the population. In *A. thaliana* and *O. sativa* TBL proteins belong to large gene families with functions related to cell wall modifications. In *A. thaliana*, At-TBL44 has been implicated in pectin esterification (Vogel *et al*., 2004; Bacete *et al*., 2018), whilst in rice other members of this family appear to be involved in acetylation of xylan moieties in cell walls (Gao *et al*., 2017). In each case, alterations in enzyme activity altered resistance in *A. thaliana* to powdery mildew and in rice to leaf blight (Vogel *et al*., 2004; Gao *et al*., 2017). Recently an 11 kDa protein was isolated from the cell wall of the shoot parasite *C. reflexa* and identified as a glycine rich protein (GRP) (Hegenauer *et al*., 2020). The protein and its minimal peptide epitope (Crip21) bind to and activate a cell surface resistance gene in tomato (CuRe1), leading to resistance to the parasite, illustrating the importance of cell wall modifications to host resistance.

In addition to cell wall modification, several candidate genes were annotated as having protease activity, including two aspartyl proteases, three subtilisin or subtilisin-like genes and a cysteine proteinase. Interestingly, all had a dual-domain predicted structure consisting of a propeptide inhibitor domain and a catalytic protease domain. In other such protease enzymes, the propeptide domain auto-inhibits the enzyme activity until cleavage of this inhibitor domain activates the catalytic domain (Shindo & Van Der Hoom 2007). This provides a mechanism by which the parasite could initially secrete an inactive VF that only becomes active once in the host environment. A similar dual-domain structure was found for a highly expressed, haustorium-specific cysteine protease in the shoot parasitic plant, *C. reflexa*, which positively contributes towards pathogenicity (Bleischwitz *et al*., 2010) Although the precise functions of other candidate VFs are unknown, for example the putative aquaporin, PAR1 protein, cytochrome P450 and the 5 proteins with no functional annotation, they provide exciting avenues for further investigation.

*S. hermonthica* has extremely high fecundity (>100,000 seeds per plant) (Parker & Riches 1993), a persistent seed bank and is obligate out-crossing (Safa *et al*., 1984), leading to a very large effective population size (Huang *et al*., 2012). Therefore, the high heterozygosity that we observed in the *S. hermonthica* genome was not unexpected. *S. hermonthica* parasitizes many different host species and varieties, often within the same geographical area. Populations therefore encounter many different forms of resistance, which they experience as a highly heterogeneous environment. This is expected to maintain genetic diversity at many loci contributing to virulence, which is consistent with observations from field studies that resistant varieties, of any particular crop species, are often parasitized by one or two *S. hermonthica* individuals (Gurney *et al*., 2006; Rodenburg *et al*., 2017). A typical example is the host-parasite combination used here as a test system; the *S. hermonthica* Kibos population and the strongly resistant upland rice variety, NERICA-17 one of 18 NERICA rice varieties grown widely by African farmers (Cissoko *et al*., 2011; Rodenburg *et al*., 2015).

This type of parasite interaction with multiple hosts leads to two predictions that are supported by our data. First, multiple loci, potentially with a wide range of functions, are likely to be implicated in overcoming host resistance. We detected 190 strong candidates for contribution to virulence, with extreme allele frequency differences between our control and virulent pools, including many gene families. It is likely that many additional candidate VFs would be revealed, by repeating this comparison on other resistant hosts. An important question for the future will be to determine how individual VFs are implicated in overcoming resistance for specific hosts or across a range of hosts. Second, maintenance of variation at virulence loci by balancing selection will lead to elevated Tajima’s D relative to the background, reflecting persistence of multiple alleles at these loci. We found the overall Tajima’s D in *S. hermonthica* to be negative, perhaps reflecting population expansion following the spread of agriculture, but our candidate loci had significantly higher Tajima’s D on average, consistent with balancing selection on these loci. Understanding the maintenance of variation at virulence loci by balancing selection will be critical to managing the evolution of virulence as a part of a sustainable control strategy (Mikaberidze *et al*., 2015).

Effective control of *S. hermonthica* is essential for food security and poverty alleviation for small-holder subsistence farmers, but it remains elusive. The use of resistance crop varieties is recognised as sustainable and cost effective (Scholes *et al*., 2008), but the durability of resistant varieties is compromised by the potential for rapid evolution of parasite virulence. Thus, the long-term success of host resistance, as a control strategy for S. *hermonthica* and other parasitic weeds, requires knowledge of the virulence factors involved, their allelic variation within and between Striga populations and their interaction with different host resistance alleles. Only then will it be possible to combine resistance alleles, in host varieties that are suitable for different agro-ecological zones and in ways that achieve sustained control by delaying the evolution of virulence. Our experimental approach and identification of candidate VFs and allelic variation within a *S. hermonthica* population, is a critical first step in this direction.

## Acknowledgements

We thank members of the library production, instrumentation and informatics teams at Edinburgh genomics. We also thank Dr Hernan Morales, University of Gothenburg, Sweden for providing the R script used to infer the read coverage distribution for each SNP for each pool of sequenced reads, based on three-component mixture models. We thank Dr Mamadou Cissoko, University of Sheffield, for help with the production of the transverse sections through rice roots infected with *S. hermonthica*. This project was funded by UKRI Biotechnology and Biological Sciences Research Council (https://bbsrc.ukri.org) grants, BB/J011703/1 and BB/P022456/1, awarded to JDS and RK and The Leverhulme Trust (https://www.leverhulme.ac.uk) grant (RPG 2013-050) awarded to JDS and RK.

## Author contributions

JDS and RKB planned and designed the research. SQ, PZ and JDS contributed to the production of *S. hermonthica* materials and extraction of DNA for genome and pooled sequencing. MB carried out library preparation and sequencing of the *S. hermonthica* genome. SQ led the genome assembly and annotation with contributions from JMB, RC, JDS and RKB. JMB carried out the prediction and analysis of the *S. hermonthica* secretome. SQ mapped the pooled *S. hermonthica* sequence reads to the *S. hermonthica* genome. SQ, RKB and JMB contributed to the population genomic analyses. JMB, PZ and JDS contributed to the analysis of changes in gene expression in *S. hermonthica* haustoria. SQ and JMB contributed equally. All authors contributed to writing of the manuscript.

## Data Availability

Raw reads for the pooled *S. hermonthica* sequences and for the *S. hermonthica* genome sequence, the assembled genome sequence and annotations have been submitted to the European Nucleotide Archive (ENA) browser at (http://www.ebi.ac.uk/ena/data/view/) under the following accession numbers: Genome Assembly GCA_902706635; Project ID PRJEB35606; Sample ID ERS4058863 and Contig accession CACSLK010000001-CACSLK010035056.

