## Supplementary figures and methods for "Identification of candidate virulence loci in *Striga hermonthica*, a devastating parasite of African cereal crops"

**Article acceptance date:** Click here to enter a date.

The following Supporting Information is available for this article:

**Fig. S1** Three step pipeline to predict the *Striga hermonthica* secretome and subsets of candidate pathogenicity-related genes.
**Fig. S2** Testing gene clustering in the secretome

**Fig. S3** Relative abundance of Pfam domains in the *Striga hermonthica* secretome or in the rest of the proteome.
**Fig. S4** Functional categorization of 4 subsets of proteins selected from the *Striga hermonthica* secretome.
**Fig. S5** Mean ΔAICcv in relation to the numbers of SNPs in 1kb windows in genic regions.
**Fig. S6** Comparison of Tajima's D for the 152 putative VFs (green) and all the genes in the genome for the control pools.
**Fig. S7** Distribution of mean ΔAICcv difference to the maximum ΔAICcv ratios in each distance interval.

**Table S1** Sequencing information for the *Striga hermonthica* reference genome and the bulked samples for pooled re-sequencing analysis (see separate file - Tables 1 – 7.xlsx).
**Table S2** Plant species included in the analysis of genome size, heterozygosity and repetitiveness (see separate file - Tables 1 – 7.xlsx).
**Table S3** Summary statistics for the *Striga hermonthica* genome assembly (see separate file - Tables 1 – 7.xlsx).
**Table S4** Repeat elements identified in the *Striga hermonthica* genome (see separate file - Tables 1 – 7.xlsx).
**Table S5** Comparison of the *Striga hermonthica* genome annotation with other plant species (see separate file - Tables 1 – 7.xlsx).
**Table S6** BUSCO completeness analysis using 2,326 core orthologous genes for eudicots (version: eudicots_odb10) (see separate file - Tables 1 – 7.xlsx).
**Table S7** Predicted subcellular location of *Striga hermonthica* proteins according to their closest ortholog in *Arabidopsis thaliana*. (See separate file - Tables 1 – 7.xlsx).

**Methods S1** – detailed list of methods and supplementary references

**Data S1** Genes encoding *Striga hermonthica* proteins (see separate xlsx file).

**Data S2** Pfam domains enriched in the secretome of *Mimulus guttatus* (see separate xlsx file).
**Data S3** FPKM values for *Striga hermonthica* haustoria during infection of the susceptible rice variety NERICA 7 (see separate xlsx file).

**Fig. S1** Three step pipeline to predict the *Striga hermonthica* secretome and subsets of candidate pathogenicity-related genes. Step 1: Three SignalP algorithms were used to identify *S. hermonthica* proteins with a predicted secretion signal at their N-terminus. These proteins were then checked for a transmembrane helix in the remaining part of the protein and only retained if no transmembrane domains were identified. Step 2: A range of structural and functional information was gathered for putative secreted proteins. Step 3: the secretome was refined into sets of candidate pathogenicity-related genes using the information gleaned in step 2. In addition to the three secretome subsets, a stand-alone BLASTp analysis was carried out against the pathogen-host interaction database.

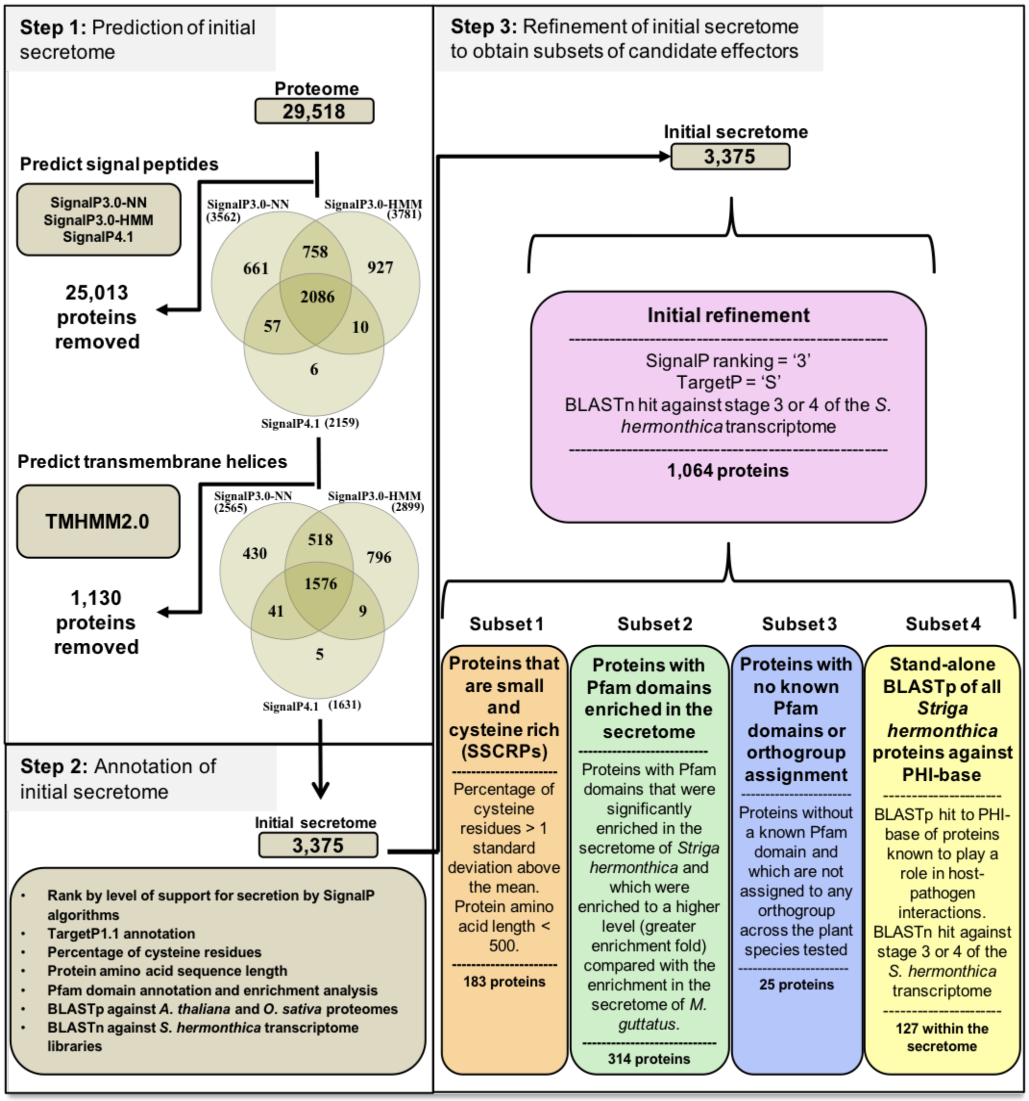

**Fig. S2 Testing gene clustering in the secretome.** The red dot represents the number of genes in the secretome with MinDis falling into each 5Kb distance interval. The upper and lower limits of the boxes are the first and third quartiles in the 10,000 permutations in each distance interval; medians of the data are shown as bands in the boxes; the whiskers represent the smallest and biggest datum that are still within 1.5 times interquartile range of the lower and upper quartile, and the outliers are shown as black dots.

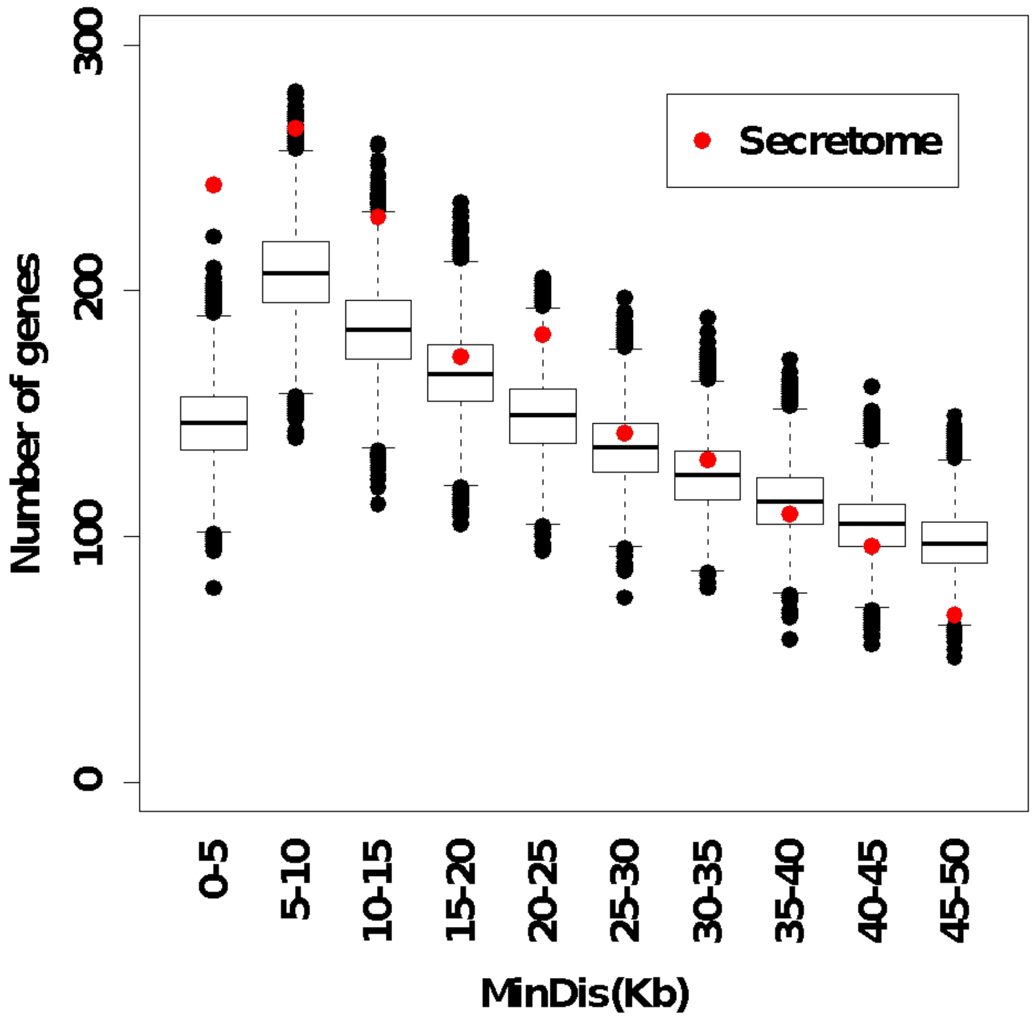

**Fig. S3 Relative abundance of Pfam domains in the *Striga hermonthica* secretome or in the rest of the proteome (non-secretome).** The relative abundance of each Pfam domain is plotted as a percentage of all Pfam domains found in the secretome or the non-secretome. The 45 Pfam domains with the highest difference in relative abundance between the two data sets are shown.

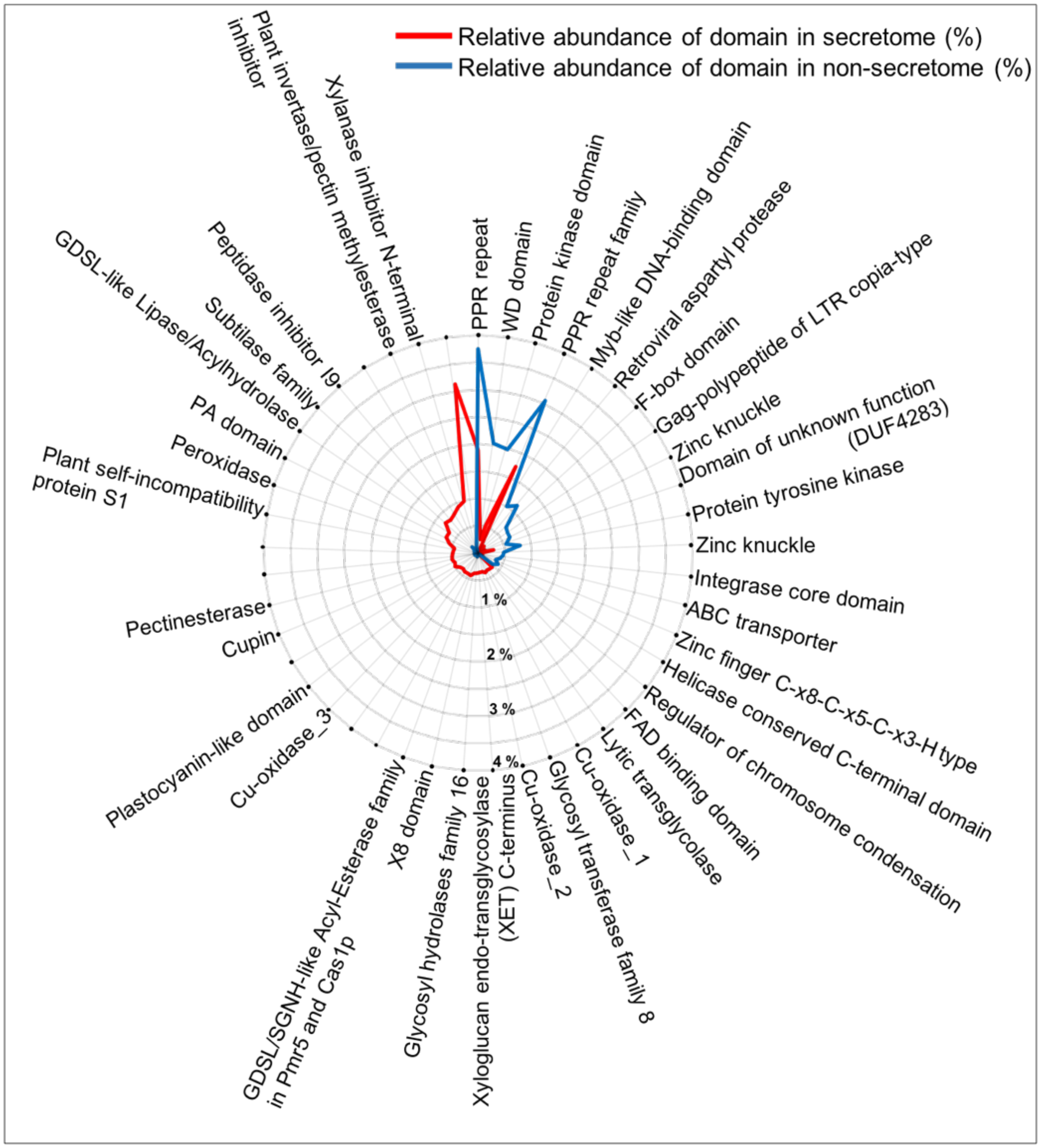


**Fig. S4 Functional categorization of 4 subsets of proteins selected from the *Striga hermonthica* secretome.** *S. hermonthica* proteins were assigned into functional groups based on the closest BLASTp match with *Arabidopsis thaliana*. If no clear annotation was available, the closest match with *Oryza sativa* was used. If no clear matches were obtained for either *A. thaliana* or *O. sativa,* the Pfam domains were used to infer function. Subset 1: small, cysteine rich proteins. Subset 2: proteins with Pfam domains enriched in the secretome of *S. hermonthica* to a greater degree than in the secretome of *Mimulus guttatus*. Subset 3: proteins with no known PFAM domain or orthogroup assignment. Subset 4: proteins with a BLASTp match against a protein in the pathogen-host interaction database. For subsets 1, 2 and 4, all *S. hermonthica* protein functions which were only represented once were categorized as 'other'. Subset 3 only contained 25 *S. hermonthica* proteins and therefore all protein functions were shown, even if they only occurred once. Venn diagram shows the overlap between the four subsets of secreted *S. hermonthica* proteins.

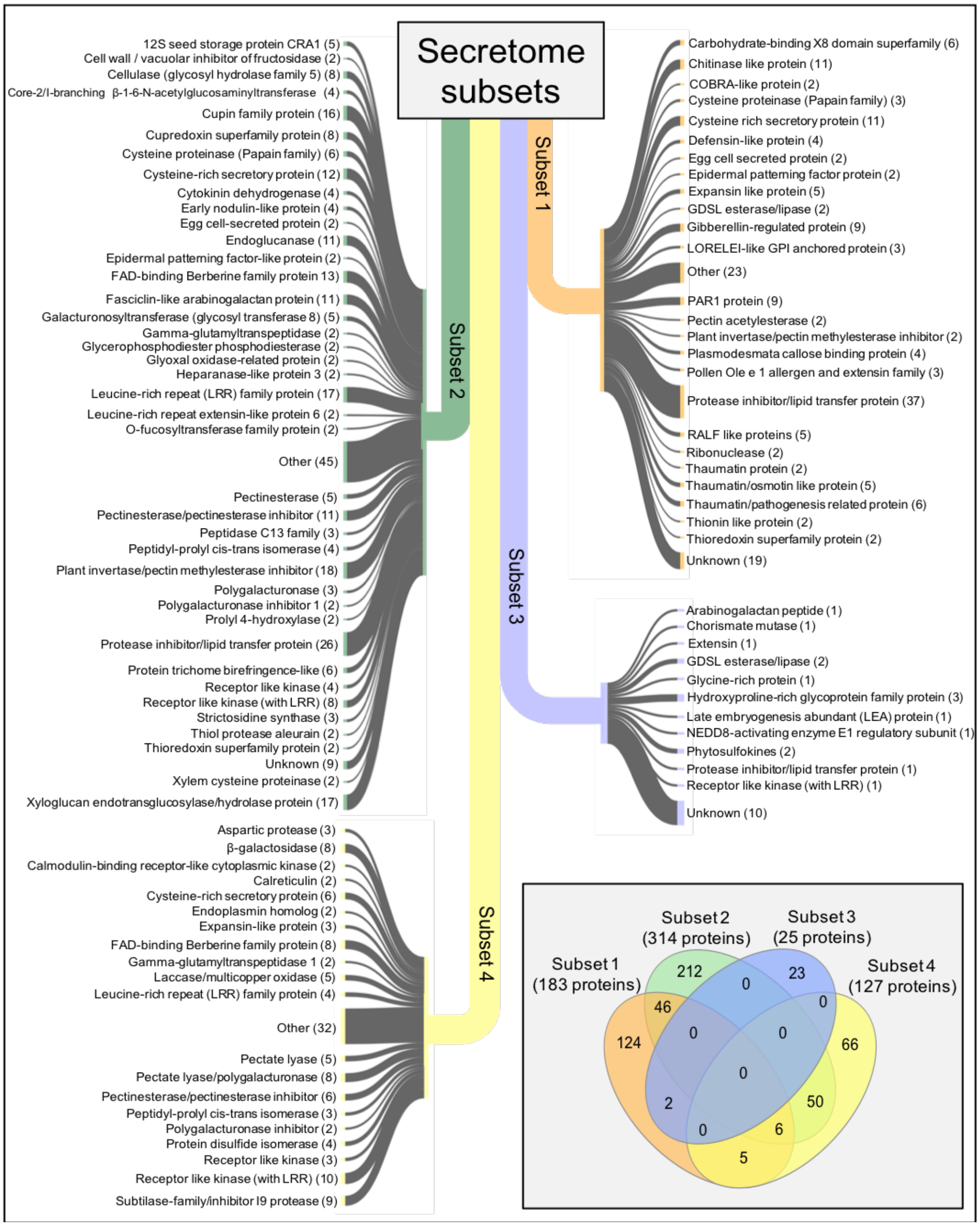

**Fig. S5 Mean ΔAICcv in relation to the numbers of SNPs in 1kb windows in genic regions.** ΔAICcv is a measure of the magnitude, and consistency across pools, of the allele frequency difference between *Striga hermonthica* growing on resistant NERICA 17 (virulent) and those growing on susceptible NERICA 7 (control). Significant windows in genic regions contributing to the secretome (red) and non-secretome (green) were identified by permutation.

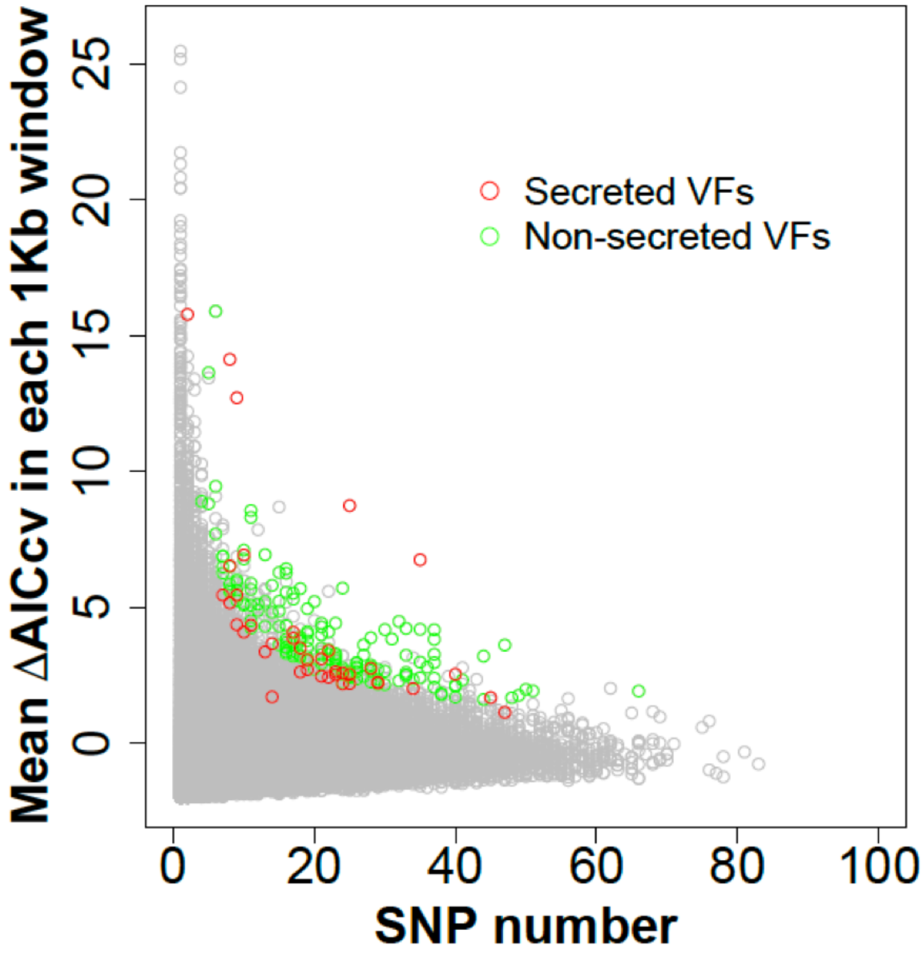

**Fig. S6 Comparison of Tajima's D for the 152 putative non secreted VFs (green) and all the genes in the genome (grey) for the control pools.** The 152 candidate loci in the proteome had significantly elevated D on average ($p < 0.0001$, $10^5$ permutations) compared to the rest of the genome.

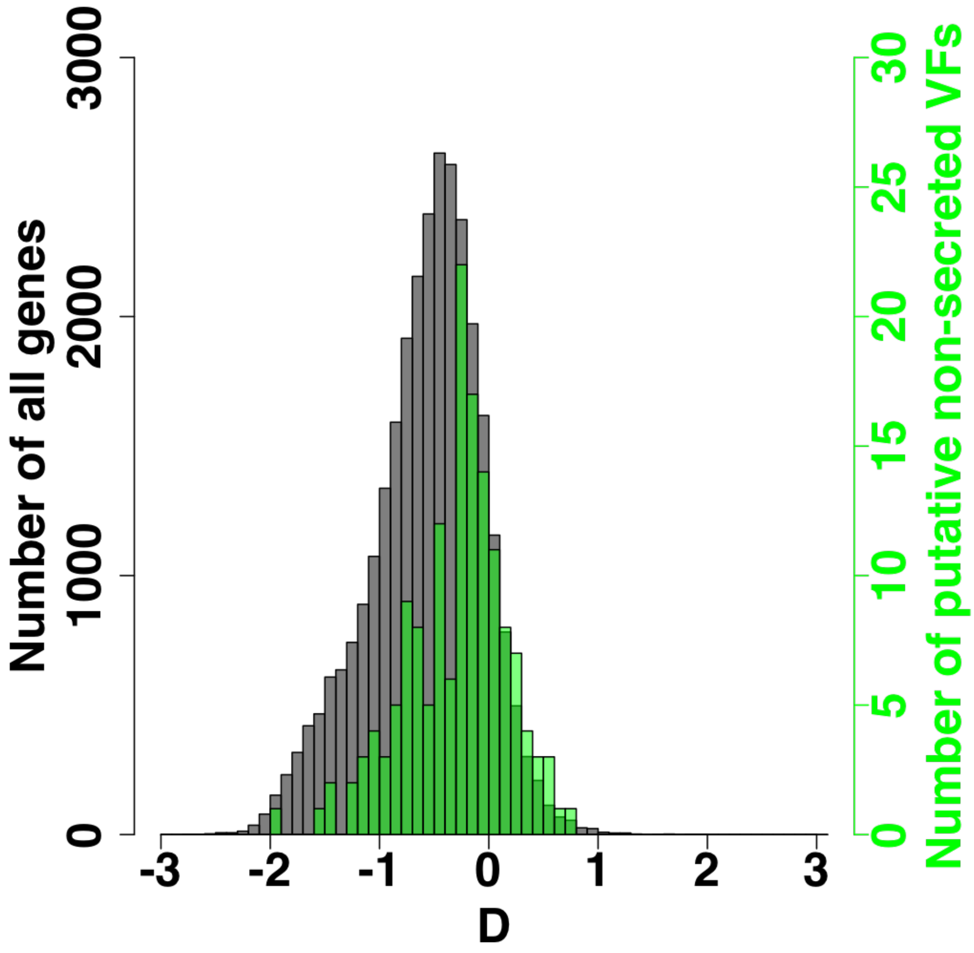

**Fig. S7 Distribution of mean ΔAICcv difference to the maximum ΔAICcv ratios in each distance interval.** The upper and lower limits of the boxes are the first and third quartiles of the ratios in each distance interval; medians of the data are shown as bands; means are shown in red dots; the whiskers represent the smallest and biggest datum that are still within 1.5 times interquartile range of the lower and upper quartile, and the outliers are shown as black dots.

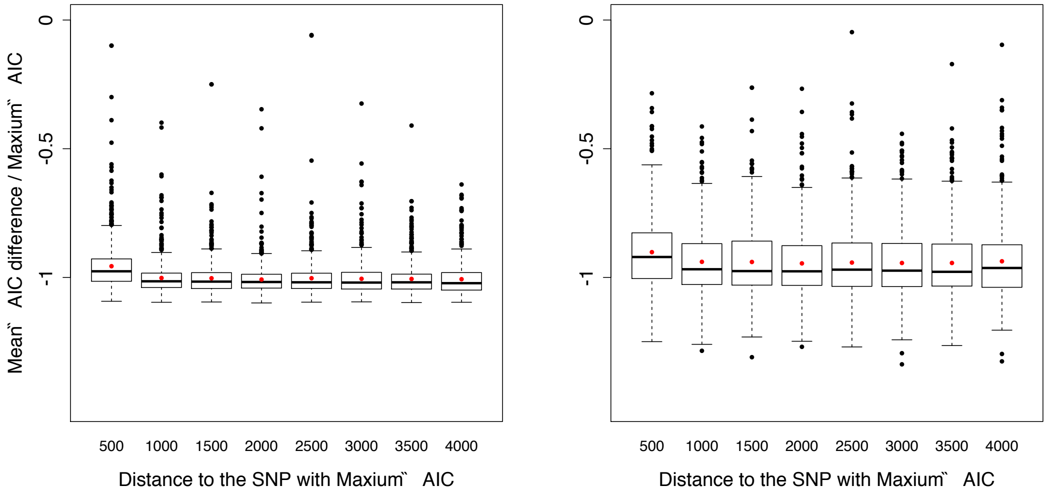


### Methods S1

#### DNA extraction from *Striga hermonthica* for genome sequencing and initial processing of sequence reads

Each pool of 100 *S. hermonthica* individuals was randomly divided into 25 groups of four individuals (20 mg of tissue per individual) for DNA extraction. To remove high levels of secondary metabolites and polysaccharides in *S. hermonthica*, lysed plant tissues were first washed in a buffer containing 2 % polyvinylpyrrolidone, 0.25 M NaCl, 0.2 M Tris-HCl and 50 mM EDTA. DNA was then extracted following the CTAB protocol (Stewart & Via, 1993). The quality of DNA was checked by gel electrophoresis and quantified by Nanodrop. An equal quantity of DNA from each of the 25 samples was then combined to form one biological replicate.

#### De novo assembly of the *Striga hermonthica* genome

For *de novo* genome assembly, adapters and low quality reads (Q < 20) were removed using CutAdapt (Martin, 2011). Only reads longer than 50 bp for the paired-end libraries and 30 bp for the mate-pair libraries were retained for subsequent analyses. Duplicated reads generated by PCR amplification in the library construction process were removed using FASTUNIQ (Xu *et al.,*

2012). Sequencing errors, which can create difficulties for the short-read assembly, were corrected using the software BLESS, a *k*-mer spectrum-based method designed to remove *k*-mers with a small number of occurrences (Heo *et al.,* 2014*).*

**Inference of orthogroups (OG)**

Orthologous gene groups (Orthogroups or OGs) among 13 plant species (*Amborella trichopoda*, *Musa acuminata*, *Oryza sativa*, *Zea mays*, *Sorghum bicolor*, *Aquilegia coerulea*, *Solanum lycopersicum*, *Mimulus guttatus*, *Striga hermonthica*, *Arabidopsis thaliana*, *Medicago truncatula*, *Populus trichocarpa* and *Vitis vinifera*) were inferred using the software OrthoFinder v2 (Emms & Kelly, 2015). Except for *S. hermonthica*, the proteomes of these species were downloaded from Phytozome V12.1 (https://phytozome.jgi.doe.gov/pz/portal.html). The number of genes per species for each OG was transformed into a matrix of *Z*-scores to quantify gene family expansion / contraction in comparison with other plant genomes. The significance of expansion or contraction was determined using CAFE v4.2 (Han *et al.,* 2013). Specifically, multi-species alignments of 42 single-copy genes from the OrthoFinder results were first constructed using MAFFT v7.407 (Katoh & Standley, 2013). A maximum likelihood (ML) tree using the alignments of the 42 single-copy genes was constructed in MEGA X (Kumar *et al.,* 2018) then converted into an ultrametric tree and calibrated based on a divergence time of 82.8 Ma–127.2 Ma between *A. thaliana* and *P. trichocarpa* (Clarke *et al.* 2011). The ultrametric tree and the gene numbers in each orthogroup from the OrthoFinder results were then used for the CAFE analysis. The functional annotation of each OG was predicted based on sequence similarity to the InterPro protein family database where the majority of the proteins in the OG shared the same Pfam annotation.

**Prediction, analysis and refinement of the *S. hermonthica* secretome**

Secreted *S. hermonthica* proteins were predicted according to the default thresholds using SignalP3.0 and SignalP4.1 (Bendtsen *et al.,* 2004; Petersen *et al.,* 2011) (Fig. S1). SignalP3.0 predicts a secretion signal peptide according to both neural network (SignalP3.0-NN) and hidden Markov model (SignalP3.0-HMM) algorithms, whilst SignalP4.1 uses a neural network algorithm. Putative secreted proteins were searched for transmembrane spanning regions in the remaining portion of the protein sequence (without secretion signal) using TMHMM2.0 (Krogh *et al.* 2001). Proteins with a secretion signal but without a predicted transmembrane helix were retained as the 'secretome'. Putative secreted proteins were ranked according to the agreement between the different SignalP algorithms: '3' = predicted by all three SignalP algorithms (SignalP3.0-NN, SignalP3.0-HMM and SignalP4.1); '2' = predicted by at least two algorithms and '1' = predicted by one algorithm. TargetP1.1 was used to provide additional predictions on the cellular localization of the putative secreted proteins ('S' = extracellular; 'C' = chloroplast; 'M' = mitochondrial) (Emanuelsson *et al.,* 2000). The closest matching *Arabidopsis thaliana* protein for each *Striga hermonthica* putative secreted protein was queried against the SUBA4 online database, which combines bioinformatic and experimental evidence to predict subcellular localisation of *A. thaliana* proteins (https://suba.live) (Hooper *et al.,* 2017). The consensus location of each queried protein was recorded and the number of each subcellular location was expressed as a percentage of the total. Using this approach, the most common subcellular location predicted in the secretome was 'extracellular', supporting the validity of our secretome prediction pipeline for *S. hermonthica* (Table S5).

To determine whether a putative secreted protein was expressed during early stages of parasitism a BLASTn search using the protein's coding sequence was conducted against the stage 3 (~ 48 hours post infection) and stage 4 (~ 72 hours post infection) transcriptome dataset downloaded from the PPGP (Lu *et al.,* 2016) BLASTn hits were taken to be significant if they returned an e-value < $10^{-10}$, percentage identity of > 95 % and a bit score of > 60. To identify enriched Pfam domains in the *S. hermonthica* secretome compared with the rest of the proteome (non-secretome), the frequency of each Pfam domain was first calculated as the number of occurrences of each domain over the total number of domains in the secretome or non-secretome. Enrichment fold for each domain was then calculated as the frequency of hits in the secretome over the frequency in the non-secretome. The significance of enrichment was assessed using a Chi-squared test with a false discovery rate (FDR) correction for multiple testing (Benjamini *et al.,* 1995) and was taken to be significant when the corrected p value was < 0.1.

The initial secretome was refined into subsets based on a series of structural and functional characteristics (Fig. S1). First, those putative secreted proteins that met the following criteria were retained: SignalP ranking = 3; TargetP location = S; and have a BLASTn hit against either stage 3 or 4 *S. hermonthica* transcriptome libraries (suggesting gene expression during early stages of parasitism). From these, four subsets of proteins were identified. Subset 1 comprised small, secreted cysteine-rich proteins (SSCRPs) $<$ 500 amino acids in length and containing $\geq$ 4 % cysteine residues. Subset 2 consisted of proteins containing Pfam domains that were significantly enriched in the *S. hermonthica* secretome. Only those Pfam domains that were enriched to a greater extent in the *S. hermonthica* secretome compared with their enrichment in the *M. guttatus* secretome (determined in exactly the same way as for *S. hermonthica*) were used to obtain this subset. Subset 3 comprised proteins with no known Pfam domains or orthogroup assignment. Subset 4 contained proteins with similarity to proteins involved in host-pathogen interactions based on having a match against the PHI-base (Fig. S1).

To test whether genes in the secretome clustered together in the genome, a permutation test was performed. For each gene, 'MinDis', defined as the distance between the start positions of the focal gene and its nearest neighbouring gene, was calculated. Genes without a neighbouring gene on the same scaffold were excluded. MinDis was also calculated for 10,000 randomly sampled gene sets with the same size as the secretome. The numbers of clustered genes with MinDis falling into a series of distance intervals were counted and were then compared with the numbers in the same distance interval in the 10,000 random gene sets.

**Identification of candidate virulence loci using pooled sequencing data**

***Trimming and filtering:*** The raw sequence reads from the six pools were trimmed to remove low quality bases and adapter sequences using CutAdapt (Martin 2011). The cleaned reads were mapped to the *S. hermonthica* reference genome using both BWA mem v 0.7.15 (Li & Durbin 2010) and NOVOALIGN (http://www.novocraft.com). Mapped reads were sorted using Samtools (Li *et al.,* 2009) and PCR duplicates were removed using Picard (http:// broadinstitute .github .io /picard/). Reads around indels were realigned using the modules RealignerTargetCreator and IndelRealigner implemented in gatk (Van der Auwera *et al.,* 2013). SNPs were called using bcftools (https://samtools. github.io/bcftools/). An R script described in (Morales *et al.,* 2019) was used to infer the read coverage distribution for each SNP for each pool of sequenced reads, based on three-component mixture models. An upper boundary of the coverage for the intermediate component (single and low copy number regions) was obtained from the fitted distribution for each pool. The high-coverage component is expected to be enriched in repetitive sequences while the low coverage component is enriched in erroneous reads or inadequately sampled genome regions. In order to provide robust data for subsequent analyses, the inference of an allele and its frequency was considered to be reliable only if the following criteria were met. (a) Coverage from each pool was $\geq 4$ and $\leq$ the upper boundary inferred from the mixture model distribution. (b) The sum of the coverage across the six DNA pools was $\geq 36$. (c) The count of the minor allele frequency, summed over pools, was > 2. (d) The ratio of the coverage of the second highest allele to the total coverage was $\geq 0.1$. (e) The ratio of the combined coverage of the third and fourth highest alleles to the total coverage was < 0.02. Only the highest and second highest frequency alleles were considered for the subsequent analyses. Although three or four alleles were present at some positions, the additional alleles were always rare. Cut-offs were determined following exploration of a range of alternative values. They removed only variants that are very unlikely to differ significantly among pools and so they did not influence the ranking of candidate loci.

The likelihood of the observed read counts for the two most common alleles, across the six pools was calculated according to equation 3 from Gompert & Buerkle (2011) to allow for the two levels of sampling associated with pooled sequencing data (sampling of reads and of individuals). We compared three allele-frequency models for each SNP using the Akaike information criterion (AIC): a null model with a single allele frequency for all pools, a control-virulent model with one frequency for the control pools (from the NERICA 7 host) and one for the virulent pools (from the NERICA 17 host) and a replicate model with a different allele frequency for each of the three pairs of pools (one control and one virulent) that were sequenced together. The control-virulent model was the model of interest while the replicate model was intended to check for consistency across pairs of pools. Therefore, two ΔAIC values were obtained: ΔAICcv = AICnull - AICcontrol-virulent and ΔAICrep = AICcontrol-virulent – AICreplicate. High positive values of ΔAICcv represent better fits than the null model and indicate significant differences between control and virulent pool types. SNPs with positive ΔAICrep values were likely to be affected by artefacts caused by sequencing methods and were excluded from the following analyses. All analysis steps were repeated independently for SNPs based on BWA and NOVOALIGN mapping as recommended by Kofler *et al.,* (2016).

***Linkage disequilibrium:*** Firstly, the distribution of the gene lengths, i.e. from the start codon to the stop codon of a gene, for all the genes in the *S. hermonthica* genome was obtained and 4 kbp, which is the 75$^{th}$ percentile of the gene length distribution, was used as the upper limit for the length of region to be considered. The SNP with maximum ΔAICcv within a gene was identified and the region up to 4 kbp from the SNP was divided into eight 500 bp intervals. The mean difference of ΔAICcv, relative to the maximum ΔAICcv, for SNPs in each 500 bp interval was then calculated. The mean difference was typically only elevated in the 500 bp interval containing the SNP with maximum ΔAICcv, so it was clear that linkage disequilibrium above background levels did not normally extend beyond 1 kbp (see Fig. S7). Therefore, 1 kbp windows were used to detect genomic regions.

***Permutation test:*** A mean ΔAICcv was calculated for a window containing n SNPs. One hundred thousand samples of n SNPs from the pool of all SNPs within the analysed regions genome-wide, were drawn randomly. For each of these samples, a mean ΔAICcv value was calculated and an expected distribution for the mean in the target window was obtained. The probability that the observed mean ΔAICcv of a window could be matched or exceeded by chance was obtained by counting the number of permuted mean ΔAICcv with values lower than the observed mean ΔAICcv value for that window and then divided by 100,000.

***Divergence measure:*** To provide a measure of divergence per gene, we used the proportion of SNPs with high $F_{ST}$. The $F_{ST}$ values between the control and virulent pools for each SNP were calculated using Popoolation2 (Kofler et al., 2011). The 95th percentile was obtained from the $F_{ST}$ values for all SNPs across the genic regions as a whole. For each genic region, we counted the number of SNPs in its significant window(s) with $F_{ST}$ value higher than the 95th percentile and then divided by the total number of SNPs in the significant window. This provided a measure of differentiation for each candidate virulence gene between the *S. hermonthica* grown on the two host varieties.

**Expression profiling of candidate virulence genes**

To determine the expression profiles for candidate virulence genes, an RNA-seq analysis was conducted for *S. hermonthica* (Kibos accession) collected at 2, 4, or 7 days post infection on NERICA 7. Rice plants were grown and infected in the rhizotron system as described previously (Gurney et al., 2006). S. *hermonthica* were collected from the roots of NERICA 7 by cutting a 1 mm section of rice root around each *S. hermonthica* attachment. In addition, unattached *S. hermonthica* haustoria were induced *in vitro* by the addition of 10 µM DMBQ (Fernández-Aparicio et al., 2013). For each treatment, four biological replicates were generated. RNA was extracted using the RNeasy plant Minikit (Qiagen) and 2 µg of RNA was treated with DNase to remove DNA contamination (dsDNase, Thermoscientific). The resulting RNA was purified and concentrated using the RNeasy MinElute cleanup kit (Qiagen) and then sent to NovoGene Co. Ltd. (China) for sequencing (https://en.novogene.com/). Cleaned reads were mapped to the *S. hermonthica* genome using Tophat2, version v2.0.12 (default settings except 'mismatch' = 2). Transcript abundance was determined as read count and measured by HTSeq (version 0.6.1) with default settings, except that the '-m' option was set to 'union'. The FPKM values for each gene at each time point were used to calculate a fold change in expression relative to the haustorial sample. To determine whether gene expression changed significantly during infection, a one-way ANOVA was carried out for each gene. If the assumptions of ANOVA were not met, data were log transformed. For each gene, log2 fold expression values, across the time points, were centred around 0 and scaled by the standard deviation for plotting as a heatmap. Heatmap plotting was conducted using the pheatmap function in R and clustering of genes by expression profiles was conducted using the 'complete' method with the 'Euclidean' distance measure.
